# Systems biology analysis using a genome-scale metabolic model shows that phosphine triggers global metabolic suppression in a resistant strain of *C. elegans*

**DOI:** 10.1101/144386

**Authors:** Li Ma, Angelo Hoi Chung Chan, Jake Hattwell, Paul R. Ebert, Horst Joachim Schirra

## Abstract

**Background:** Pest insects are increasingly resistant to phosphine gas, which is used globally to protect grain reserves. The enzyme dihydrolipoamide dehydrogenase (DLD) is a phosphine resistance factor and participates in four key steps of core metabolism, making it a potential central metabolic regulator.

**Results:** Here we used microarray data and NMR-based metabolomics to characterize the phosphine response of wild-type *C. elegans* and the phosphine-resistant strain *dld-1*(*wr4*) which has a partial loss-of-function mutation in the gene for DLD. In addition, we have constructed *Ce*Con, a *C. elegans* genome-scale metabolic model to facilitate integration of gene expression and metabolomics data.

**Conclusions:** The resulting systems biology analysis is consistent with the hypothesis that adaptation to a hypometabolic state is the most prominent mechanism of phosphine resistance in this nematode strain. The involvement of DLD in regulating and creating hypometabolic adaptation has implications for other biological phenomena involving hypometabolism, such as reperfusion injury and metabolic resistance.

## Background (words 1561)

Phosphine (PH_3_) is the sole fumigant registered for the routine protection of stored grain from insect pests. It is an effective metabolic toxin that kills actively aerobically respiring organisms, but which spares grain as grain is metabolically dormant. However, multiple insect species that are grain storage pests have developed a high level of resistance against phosphine in countries across the world (Collins et al. 2002). The continued use of this very important chemical will depend on an understanding of the mechanisms of toxicity and resistance.

Recently we have shown that variants of dihydrolipoamide dehydrogenase (DLD), a key enzyme of energy metabolism, are responsible for high-level phosphine resistance in two pest insect species, as well as in the model organism *C. elegans* (Schlipalius et al. 2012). As the metabolic mode of action of phosphine is unknown, the metabolic adaptation mechanisms that might lead to resistance are likewise unclear. The availability of a phosphine resistance mutation in the *dld-1* gene of *C. elegans* provides an opportunity to exploit the excellent genome annotation of *C. elegans* to understand phosphine toxicity and resistance, by comparing both the metabolic and gene expression responses to phosphine exposure between resistant and wild type animals.

Evidence that phosphine is a respiratory poison includes inhibition of respiration upon exposure to phosphine. For example, phosphine inhibits oxygen consumption in isolated mitochondria from rat liver (Nakakita et al. 1971) and insects (Chefurka et al. 1976). It also inhibits oxygen consumption in intact insects (Bond et al. 1969) and nematodes (Zuryn et al. 2008). In addition, ATP levels decrease in insects (Price and Walter 1987) and both ATP levels and mitochondrial membrane potential have been shown to decrease in nematodes (Valmas et al. 2008, Zuryn et al. 2008) following exposure to phosphine.

Cytochrome c oxidase has been proposed to be the direct target of phosphine as the activity of cytochrome c oxidase is inhibited when isolated mitochondria are exposed to phosphine gas (Chefurka et al. 1976). Inhibition is not observed, however, when live insects are exposed to phosphine gas, followed by isolation and analysis of their mitochondria (Price 1980, Price and Dance 1983). Interestingly, cytochrome c oxidase activity of rat brain or human blood is inhibited by almost 50% after *in vivo* exposure to aluminium phosphide in solution (Dua and Gill 2004, Singh et al. 2006). Interpretation of the mammalian result is confounded by the toxicity of aluminium to the electron transport chain (Halliwell 1992) as well as the strong inhibitory effect of aluminium at micromolar concentrations to transcription of cytochrome oxidase subunit 3 (Bosetti et al. 2001).

An alternative explanation of phosphine toxicity is that, rather than causing lethal metabolic insufficiency, mitochondrial inhibition results in the generation of reactive oxygen species (ROS). The ROS are then proposed to damage the macromolecules of the cells resulting in lethal oxidative stress. Observations supporting this hypothesis included increased H_2_O_2_ levels in phosphine treated mitochondria from insects (Bolter and Chefurka 1990), as well as increased ROS-associated oxidative damage in insects (Chaudhry and Price 1992), nematodes (Valmas et al. 2008), rat tissue (Dua and Gill 2001, Hsu et al. 2000, Hsu et al. 2002b) and mammalian cell lines (Hsu et al. 1998) following exposure to phosphine. Furthermore, synergistic enhancement of phosphine toxicity is observed when oxygen levels are elevated during fumigation (Bond 1963, Cheng et al. 2003).

Despite a large body of circumstantial evidence that links phosphine to oxidative damage, there is no definitive evidence that oxidative stress is responsible for the lethality of phosphine. For example, reports from Hsu (Hsu et al. 2002a, Hsu et al. 2000) demonstrated that antioxidants could reduce phosphine-induced oxidative damage, but they never showed that this improved the survival rate of animals exposed to phosphine. Others have shown that depletion of cellular glutathione enhances the toxicity of phosphine in mammals (Hsu et al. 2002b) as well as nematodes (Valmas and Ebert 2006), but this does not demonstrate that the normal endogenous glutathione levels are not adequately protective. In fact, phosphine exposure has been shown to result in an increase in cellular glutathione (Shivanandappa and Rajendran 1987). There is convincing evidence that catalase and peroxidases actually fail to provide protection against phosphine toxicity, despite their key role in protecting cells from ROS (Price and Walter 1987). Despite this contrary evidence, damage due to ROS remains the favoured mechanism to explain the toxicity of phosphine.

The two previous hypotheses presume that the toxicity of phosphine results from metabolic dysfunction leading to either metabolic insufficiency or the generation of ROS. There is also evidence for a third type of interaction between metabolism and phosphine toxicity. Chronic suppression of metabolism is protective, whereas an increased metabolic rate increases the toxicity of phosphine (Schlipalius et al. 2006). Examples of protective metabolic suppression include such conditions as cool temperature, genetic suppression of the mitochondrial electron transport chain (Zuryn et al. 2008) and hypoxia (Bond 1967, Kashi 1981). We have also found that the metabolic rate of the phosphine resistant mutant of *C. elegans dld-1*(wr4) is suppressed to about 25% of normal levels (Zuryn et al. 2008). In contrast, increasing metabolic demand via the administration of uncouplers dramatically enhances the toxicity of phosphine (Valmas et al. 2008).

The positive relationship between metabolic rate and the toxicity of phosphine is completely consistent with the observation that dormant organisms (e.g. seeds) are extremely resistant to phosphine. Hypometabolism would essentially allow the effect of a metabolic poison to be avoided (Schlipalius et al. 2006). Adaptation to a hypometabolic state is achieved through metabolic restructuring that sets the balance between energy demand and energy supply to a significantly lower level. Examples of hypometabolism include that induced by extreme anoxia in some aquatic turtles as well as hibernation in mammals, diapause in insects and nematodes and hydrogen sulfide induced suspended animation. Interestingly, the *dld-1*(*wr4*) mutation that causes resistance to phosphine also causes a constitutively hypometabolic state (Zuryn et al. 2008).

Given the competing theories outlined above, systems biology provides a logical approach to characterise phosphine toxicity and elucidate the mechanism of phosphine resistance. This involves monitoring biological responses with transcriptomics and metabolomics together with with bioinformatic and computational analyses to yield a better understanding of these complex biological processes. *C. elegans* provides an ideal model organism for this research, as it can be cultured under environmentally controlled conditions and it is possible to analyse 1,000s of individuals per condition, making inter-sample differences small. In addition, it is a strongly inbreeding organism, which leads to essentially complete homozygosity of individuals as well as genetic uniformity between strains. For example, only two non-synonymous nucleotide variants other than the dld-1(wr4) mutation itself distinguish the *dld-1* mutant from the N2 control strain). Available genetic resources include 10s of thousands of well-characterized near isogenic mutant strains of *C. elegans* as well as *E. coli* strains that can be fed to *C. elegans,* resulting in epigenetic suppression of almost any *C. elegans* gene.

Mapping of metabolic pathways that exist in an organism provides a biologically rich foundation for the analysis of -omics data, such as genome-wide gene expression and metabolite profiles. This approach enables holistic visual analysis that facilitates identification of biological processes that may be obscured in a purely statistical analysis. Despite the value of applying metabolic models to the analysis of -omics data, it is still more common to rely on quantitative statistical approaches.

The goal of the statistical strategy is to identify biological processes, typically represented by gene ontology (GO) terms, whose associated genes are disproportionately represented among the subset of genes that are differentially regulated in response to a stimulus. This approach may miss processes with one or a few genes that are differentially regulated. Thus, it is inherently biased against metabolic pathways that may be regulated by gene expression changes in one or two rate-limiting steps. In contrast, structures such as ribosomes that rely equally on many component subunits for their function are much more likely to be reliably identified by quantitative statistical analysis. Thus, it is clear that metabolic models enhance the insight that can be obtained from a purely quantitative statistical analysis of genome-wide data.

Two major approaches to mapping metabolic pathways have been developed that have been built upon fundamentally different philosophical bases. One of these is the Kyoto encyclopedia of genes and genomes (KEGG), which relies on a composite metabolic map derived from all organisms. The other approach is represented by genome-scale metabolic models (GSMs), as developed by Palsson [REF] and incorporated into the MetaCyc (metabolic encyclopedia) family of species-specific metabolic pathway maps. The curated, species-specific GSMs are rich in information that is relevant to the target species, whereas the composite metabolic model that forms the basis of KEGG includes a great deal of information that is simply not applicable to each specific species.

Using quality annotation data from the UniProt and Wormbase databases, we have generated *Ce*Con (*<u>C</u>. <u>e</u>legans* metabolic re<u>Con</u>struction), a GSM of *C. elegans,* on which we then mapped both experimental gene expression data as well as metabolomics data. With the same data we also carried out a quantitative statistical analysis of metabolic gene expression changes and of changes in metabolite levels for comparative purposes. This combined approach has allowed us to evaluate several metabolic interpretations of *dld-1* mediated phosphine resistance in *C. elegans.* We present the constitutive differences between strains, as well as the differing responses of resistant and susceptible *C. elegans* to phosphine exposure.

## Methods

### Nematode strains and culture conditions

The phosphine resistant *C. elegans* mutant, *dld-1*(*wr4*), was derived from the wild type strain, *N2* (var.Bristol) (Cheng et al., 2003) and was outcrossed to the *N2* strain 5x. All nematodes were cultured at 20°C on NGM agar (0.3% NaCl, 0.25% peptone, 5mg/ml cholesterol, 1mM CaCl_2_, 1mM MgSO_4_, 1.7% agar) seeded with *Escherichia coli* (OP50).

Synchronised cultures of nematodes were obtained by exposing gravid females to alkaline sodium hypochlorite solution to isolate their eggs (Stiernagle, 2006). The eggs were allowed to hatch overnight in aerated M9 buffer, which does not support further growth upon hatching. The young nematodes were then transferred to NGM agar plates that had been seeded with OP50 to initiate synchronized growth. After growth for 48 hours at 20 °C, the resulting synchronised young adult nematodes were exposed to phosphine.

### Phosphine exposure

The LC50 for adults of the *dld-1* mutant strain toward phosphine is 4000 ppm, whereas that of N2 is 500 ppm when exposed to phosphine for 24 hours at 20°C. For RNA extraction for microarray analysis, synchronised young adult nematodes of the N2 strain were exposed to phosphine at 0 ppm (air control) or 500 ppm (24 hour LC50 for N2) in sealed desiccator chambers for 5 hours at 20-21°C. Mutant *dld-1*(*wr4*) nematodes were exposed to 0 ppm, 500 ppm (sublethal for *dld-1*(*wr4*)) or 4000 ppm phosphine (24 hour LC_50_ for *dld-1*(*wr4*)) for 5 hours at 20-21°C.

The same conditions were used to prepare worms for metabolite extraction, except that the exposure time was increased to 6 hours to allow time for changes in gene expression to be reflected in the metabolite levels. In addition, the N2 strain was also exposed to a sublethal concentration of phosphine, 70 ppm, to correspond to the sublethal 500 ppm exposure for the *dld-1*(*wr4*) mutant strain.

### Microarray hybridisation

After 5 hours exposure to phosphine, worms were transferred to microcentrifuge tubes immediately and washed 3 times with M9 buffer. RNA was isolated using a Qiagen RNeasy^®^ Mini Kit. RNA samples were purified from 3 independent biological replicates for each of the 5 treatment conditions (*N2* air, *N2* 500 ppm, *dld-1* air, *dld-1* 500 ppm, *dld-1* 4000 ppm). After isolation, the RNA was sent to the Institute of Molecular Bioscience (IMB) Microarray Facility (Brisbane, Australia) for hybridization. The cRNA was synthesized from 500 ng of total RNA by a One-Color Aglient Quick Amp Labeling Kit. cRNA was fragmented and then hybridized to the Aglient *C. elegans* (V2) Gene Expression Microarray Genechip that contained 4x 44K gene probes covering approximately 2,000 *C. elegans* genes by the IMB microarray facility.

### Microarray data analysis

Microarray data analysis was performed using GeneSpring GX 10.0.2 software. Probe duplicates on the array were averaged to a single value before analysis, but multiple unique probes were analysed independently. We used the quantile normalization algorithm for data normalization among the 15 gene expression profiles.

Three expression ratios were calculated from this data:

1. The *N2* lethal response – the change in gene expression levels of *N2* when exposed to 500ppm phosphine, compared to 0 ppm phosphine
2. The *dld-1* sub-lethal response – the change in gene expression levels of *dld-1* when exposed to 500ppm phosphine, compared to 0 ppm phosphine
3. The *dld-1* lethal response – the change in gene expression levels of *dld-1* when exposed to 4000ppm phosphine, compared to 0 ppm phosphine

Subsequently, the expression ratios were used to derive three filtered expression ratios columns:

1. The survival-specific response – the *dld-1* sub-lethal response values were only retained if the changes in the two lethal responses was below the twofold increase or decrease threshold.
2. The fatality-specific *N2* response – the *N2* lethal response values were only retained if the response in *dld-1* sub-lethal was below the twofold increase or decrease threshold.
3. The fatality-specific *dld-1* response – the *dld-1* lethal response values were only retained if the response in *dld-1* sub-lethal was below the twofold increase or decrease threshold.

### Genes whose expression were differentially regulated between strains in response to phosphine exposure

Two-way ANOVA (*y_ij_* = α_*i*_ + β_*j*_ + [αβ]_*ij*_ + ε_*ij*_) was used to identify significant gene expression differences between the phosphine sensitive *N2* and phosphine resistant *dld-1* nematode strains. The first comparison examined gene expression between the two strains under equal exposure to phosphine (500 ppm), whereas the second made the comparison under equivalent physiological stress (i.e. the respective LC_50_ of each strain). In the first analysis, four exposure conditions were compared: *N2* exposed to air, *N2* exposed to 500 ppm phosphine, *dld-1* exposed to air, *dld-1* exposed to 500 ppm phosphine. In this case, α is the main effect for strain, β is the main effect for exposure, αβ is the interaction effect between strain and exposure. In the second 2-way ANOVA model the four exposure conditions that were compared were: *N2* exposed to air, *N2* exposed to 500 ppm phosphine, *dld-1* exposed to air, *dld-1* exposed to 4000 ppm phosphine. In this case, α is the main effect for strain, β is the main effect for physiological stress (LC50), αβ is the interaction effect between strain and physiological stress. For the purpose of identifying potential resistance mechanisms, we were particularly interested in genes with an interaction effect in either of these two 2-way ANOVA analyses.

To reduce the number of falsely identified interactions, the Benjamini and Hochberg multiple testing correction (MTC) algorithm (Benjamini and Hochberg 1995) was set to a false discovery rate of 0.05. To limit the identified genes to those most likely to be responsible for biologically significant effects, we also used a relative magnitude of change filter. Firstly we required that induction or suppression of gene expression due to phosphine exposure was at least 2 fold in at least one of the strains. Furthermore, we required that the magnitude of gene expression change in the other strain was less than half that in the original strain if regulation occurred in the same direction. This generated four gene subsets:

1. genes induced at least 2 fold in *dld-1*(*wr4*), but either suppressed in N2 or, if induced, induced by less than half that of *dld-1*(*wr4*).
2. genes suppressed at least 2 fold in *dld-1*(*wr4*), but either induced in N2 or, if suppressed, suppressed by less than half that of *dld-1*(*wr4*).
3. genes induced at least 2 fold in N2, but either suppressed in *dld-1*(*wr4*) or, if induced, induced by less than half that of N2.
4. genes suppressed at least 2 fold in N2, but either induced in *dld-1*(*wr4*) or, if suppressed, suppressed by less than half that of N2.

These four gene subsets were then grouped according to available annotations and enrichment scores were determined using the online Database for Annotation, Visualization and Integrated Discovery (DAVID) (Dennis et al.,2003). This directed us to several major biological processes that were overrepresented in one of the four gene subsets. The results related to the metabolic model are discussed in this paper.

Multiple testing correction is based on the assumption that each comparison is independent of all others. The reality, however, is that the gene expression responses to extreme stress is necessarily highly coordinated, particularly genes that are involved in the same biological process or metabolic pathway. Therefore, the number of independent gene expression responses is likely less than the total number of genes, suggesting that the multiple testing correction is likely to be overly stringent and cause significant expression changes to be missed. We dealt with this problem by initially using genes that passed the multiple testing correction to identify biological processes whose genes were overrepresented in one of the four gene subsets. Then additional genes were included in the pathways if they passed the interaction P-value 0.05 without the MTC being applied. The two-fold change filtering criterion was used in all cases.

### Genes showing expression differences between strains when exposed to only air

Gene expression differences that exist between wild type and phosphine resistant strains and that contribute to the resistance phenotype need not be responsive to phosphine. Therefore, we used a t-test to identify genes whose expression differed constitutively between the two strains. We used the Benjamini and Hochberg multiple testing correction algorithm (Benjamini and Hochberg 1995) to limit false positive gene identification to a frequency of 0.05. We also required that gene expression differ by at least 2 fold.

### Data presentation

The pathways identified by the quantitative analysis have also been displayed in the context of metabolic pathways to facilitate comparison with the results emanating from the metabolic model. The data are plotted as the log2 expression values for each strain at 0, 500 and 4000 ppm phosphine. To facilitate comparison between genes, the expression values in the plots have been normalized to the average of the five expression values for each gene. Error bars represent the standard error of the means across three biological replicates. In the figures, gene expression data that passed the multiple testing correction are boxed in red. Data that passed the initial statistical threshold, but not the multiple testing correction are included for the reasons discussed above, but are not highlighted in red. Data from genes that contribute non-sequentially to a process such as a stress response or that are members of a gene family are presented in tables.

### Metabolomics nematode extraction

After 6 hours exposure to phosphine, ~2000 nematodes per NMR sample (*n*≥6 for each condition) were transferred to sterile polypropylene tubes by washing the NGM plates twice with 5 mL ice-cold M9 buffer (Strange et al. 2007). From wash until the addition of solvent for extraction, samples were kept on ice to prevent metabolic activity. After 1 min centrifugation at 33 rpm and 4°C, the supernatant was removed, until ~2 mL remained. After repeated centrifugation at 33 rpm and 4°C for 1 min the remaining supernatant was removed. The nematode pellet together with residual buffer was transferred into tared microcentrifuge tubes and metabolites were extracted with a methanol-water extraction (Beckonert et al. 2007). 4 mL/g ice-cold methanol were added to the nematode pellets and the nematodes were homogenised with a Qiagen TissueLyser II ball mill at 30 Hz for 8 minutes with three 3 mm round glass beads in every tube. The resulting suspension was centrifuged at 12,000 *×g* for 10 min at 4°C. The supernatant was collected, and the pellet was re-extracted in a second round of extraction with ice-cold 80% (v/v) methanol. Supernatants for each sample were combined and dried overnight in a centrifugal vacuum evaporator. In total 39 extracts were prepared (Supplementary Table 1).

### NMR spectroscopy

The dried nematode extracts were resuspended in 200 μL 100 mM sodium phosphate buffer, pH 7.4 in 90% D_2_O containing 100 μM 2,2-dimethylsilapentane-5-sulfonic acid (DSS) as chemical shift reference and 100 mM 1,1-difluoro-1-trimethylsilanylphosphonic acid as pH control. 1D-NOESY spectra were measured at 298 K using a chilled automatic sample changer on a Bruker AV900 NMR spectrometer (Bruker Biospin), equipped with a 5 mm z-gradient triple resonance probe. 256 scans were acquired at 32k resolution over 14 ppm spectral width. Water suppression was achieved by low-power continuous-wave irradiation on the residual water resonance during the 3 s relaxation delay and 0.1 s NOESY mixing time. In addition 1D Carr-Purcell-Meiboom-Gill (CPMG) spectra were measured to suppress signals from large molecular weight constituents. 256 scans were acquired at 32k resolution over 14 ppm spectral width. The CPMG sequence used an interpulse delay of 500 μs and a loop counter of 400, leading to a total spin-lock time of 200 ms. The residual water signal was suppressed by low-power continuous-wave irradiation during the 3 s relaxation delay. For metabolite identification, 2D-TOCSY, ^13^C-HSQC, ^13^C-HSQC-TOCSY, and ^13^C-HMBC spectra were acquired as previously described (Li et al. 2011, Schirra et al. 2008).

### Processing and data reduction

Raw spectra were multiplied by a sine function phase-shifted by π/2, Fourier-transformed, phase/baseline-corrected, and the chemical shift referenced to DSS (0 ppm). NMR signals in all spectra were aligned with each other in MATLAB with the icoshift routine (Savorani et al 2010). Afterwards, spectra were data-reduced with an in-house MATLAB script between 10.0-0.3 ppm into regions (“buckets”) of 0.01 ppm or 0.001 ppm width by integrating signal intensity and normalizing to total spectrum intensity. The regions 5.01-4.66 and 3.36-3.33 ppm containing the signals of water and methanol, respectively, were excluded before normalization.

### Targeted metabolite profiling

The normalized relative concentrations of identified metabolites were calculated by summing up intensities for specific, non-overlapping signals in the data-reduced spectra. Fold-changes were calculated by dividing the relative concentrations under two different exposure conditions by each other. This yielded fold-changes for the lethal and sub-lethal responses of the wild-type and *dld-1* (mutant) nematodes similar to the microarray data analysis described above. In addition, the normalized relative metabolite concentrations were analysed by 2-way ANOVA and visualized as log2 transformed data as described above for the microarray data.

### Generating a *C. elegans* genome-scale metabolic model

A *C. elegans* GSM was generated using *C. elegans* specific gene annotations obtained from UniProt and Wormbase. These annotations were supplemented with gene annotations for orthologous genes of other well-annotated organisms in the *Ensembl* database, namely *H. sapiens, M. musculus, D. melanogaster, S. cerevisiae* and *E. coli.* Probable orthologous proteins were identified as reciprocal best matches using a proteome x proteome BLASTp search. To obtain multispecies-normalized alignment scores for each putative orthologue, the score for the reverse alignment was used (i.e. gene from other organism as query against *C. elegans*). As all scores were derived from searches using a single protein query against the *C. elegans* proteome, scores were not distorted by differences in the sizes of the proteome of each species. Annotations from orthologous proteins were only used if they were consistent between the multicellular eukaryotes. *S. cerevisiae* and *E. coli* annotations were only used to provide additional insight.

These annotated proteins were matched against the Biocyc database of metabolic pathways, using Pathway Tools software, and assembled into a GSM for *C. elegans.* Pathway Tools uses a combination of EC numbers, enzyme names, and GO terms to match input enzymes to its knowledgebase when constructing the GSM. Transporters and protein complexes were manually specified using the Pathway Tools interface. A mitochondria-specific metabolic pathway map and a lysosome-specific metabolic pathway map were also constructed.

A detailed instruction manual for generating and using CeCon, complete with examples is available in the supplementary materials. *Ce*Con is accessible online at *http://cai-gsmodel.cai.uq.edu.au*.

### Comparison of CeCon to published genome scale models

To compare the coverage and size of our GSM to the published models, ElegCyc (Gebauer et al. 2016) and iCEL1273 (Yilmaz & Walhout 2016), copies of the two models were obtained. ElegCyc was obtained in Pathway Tools format from the authors, and iCEL1273 was downloaded in Microsoft Excel format from the WormFlux website (*http://wormflux.umassmed.edu*). The models were compared in a variety of metrics including the number of pathways, reactions, enzymes and compounds, as well as compartments included in the model. As both our metabolic pathway map and ElegCyc were developed in Pathway Tools (version 19.5 and 16 respectively), statistics were obtained from the software. To compare to iCEL1273, the values were obtained from a combination of the WormFlux website, the iCEL1273 publication (Yilmaz & Walhout 2016) and the model file itself. In addition, an unpublished, but publicly available metabolic reconstruction of *C. elegans* was also used for this comparison.

The comparative size of the models was determined manually, due to the differences in software and formatting. This was done in a pathway-by-pathway manner, in which pathway lists for each model were created and combined into a “super-list”. Each model was examined for the pathways in the super-list and the data tabulated. This pathway approach was chosen as it provides a clear overview of the contents of the metabolic models. Where similar pathways existed between models, the pathways were grouped. For example, each model organized β-oxidation of fatty acids in a different fashion, so the three pathways were grouped under Fatty Acid Oxidation. Finally, using the MetaNetX namespace, the models were compared using the reaction lists.

### Overlaying experimental microarray and metabolomics data on the metabolic pathway map

The GSM facilitates the qualitative visual analysis of genome-wide metabolic data. This is achieved by colour-coding both the magnitude and direction of changes in gene expression or metabolite levels and overlaying this information onto the metabolic pathways map using the Pathway Tools software, resulting in presentation of complex datasets in the functional context of metabolic pathways. As a results, anomalies and inconsistencies are readily distinguished from coordinated and directionally consistent changes in gene expression. Rather than apply statistical filters to the data, every gene or metabolite that could be assigned to a metabolic pathway or transporter was included in the visual analysis with the magnitude and direction of gene expression changes indicated by a colour code.

## Results

To understand how organisms develop resistance to phosphine, we carried out genome-wide gene expression profiling as well as NMR-based metabolomics on *C. elegans* of the wild type *N2* strain in comparison with the phosphine resistant strain, *dld-1*(*wr4*). By observing strain x treatment interactions as well as constitutive differences in gene expression or metabolite levels between strains, we have been able to identify genes and metabolic pathways whose regulation is correlated with resistance as well as constitutive differences between strains that may also contribute to resistance. We have also been able to identify current theories of resistance that are not supported by our data.

### Statistical analysis of the microarray and metabolomic data

The initial inter strain comparison of the gene expression response to phosphine exposure was carried out at 500 ppm phosphine. This level of exposure kills 50% of *N2* if maintained for 24 hours at 20°C (LC_50_), but it is essentially non-lethal to *dld-1*(*wr4*) under the same conditions. To prevent the confounding effect of death on the experimental results, exposures were terminated at 5 hours, well before phosphine induced lethality was triggered. We used two-way ANOVA to identify genes or metabolites whose regulation is influenced by interaction between two factors, strain (*N2* versus *dld-1*(*wr4*)) and phosphine exposure, either at the same dose (500 ppm) (Figure 1A) or at concentrations that resulted in the LC_50_ of each strain (500 ppm for N2 versus 4000 ppm for *dld-1*(*wr4*)) (Figure 1B). In figure 1 and throughout the text, the term “dose” indicates a comparison in which both strains were exposed to 500 ppm phosphine and “stress” indicates a comparison in which both strains were exposed at their respective LC_50_s.

**Figure 1:**
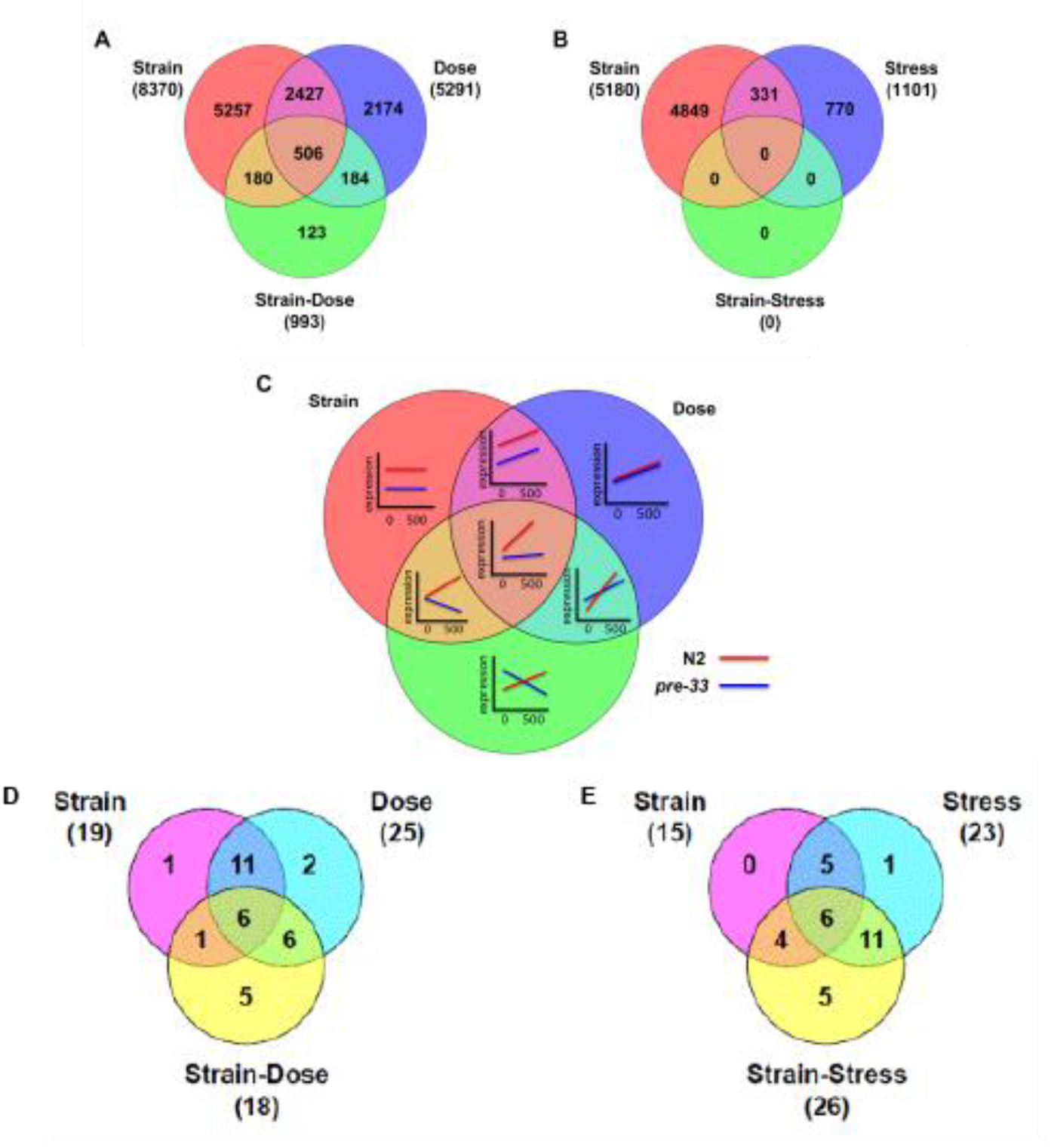
Venn diagrams showing the number of genes significant in A) chemically equivalent doses (0 and 500ppm) and B) doses which produce equivalent stresses (0, and the respective LC_50_). Panel C illustrates response types that are characteristic of each sector of the Venn diagrams. Panels D and E show the number of metabolites that are significant in the same comparisons as panels A and B.

The “dose” analysis summarised in figure 1A was used to identify genes or metabolites whose expression in response to phosphine was specifically associated with the resistance phenotype of the *dld-1*(*wr4*) strain. Specifically, we selected genes/metabolites for which responses differed between strains in response to 500 ppm phosphine. The diagram in figure 1C illustrates the types of responses that are characteristic of each sector of the Venn diagrams. The full range of expression/concentration patterns can be viewed in supplemental data (Figure s3-6). Characteristic patterns of gene expression or metabolite concentration changes that represent an interaction between strain and 500 ppm exposure are We reasoned that the responses shown in the 4 lowest sectors (i.e. the green circle) of figure 1C would be characteristic of genes contributing to the resistance phenotype of *dld-1*(*wr4*).

We refined the group of candidate resistance response genes further by applying two-way ANOVA to the two parameters, strain and exposure to 500 ppm phosphine. We also applied the Benjamini and Hochberg multiple testing correction at a false discovery rate of 0.05, resulting in a statistically significant subset of 993 interacting genes and 27 metabolites from those displayed in figure 1A and Figure 1D. These 993 genes had already passed the magnitude change filter that was applied in the initial analysis as described in materials and methods. The genes and metabolites fell into four response categories as follows:

1. 203 Group 1 - induced in *dld-1*(*wr4*)
2. 332 Group 2 - suppressed in *dld-1*(*wr4*)
3. 189 Group 3 - induced in *N2*)
4. 17 Group 4 - suppressed in *N2*)

Genes in group 1 and group 2, which responded more strongly in *dld-1*(*wr4*) than in N2 are potentially responsible for the resistance of *dld-1*(*wr4*) to 500 ppm phosphine. Genes in group 3 and group 4, which responded more strongly in *N2* than in *dld-1* at 500 ppm exposure, may have contributed to the mortality of *N2* at this dose.

The “strain” analysis summarised in figure 1B was used to identify genes or metabolites whose expression response to phosphine differed between strains despite an equivalent level of induced mortality. Such genes would constitute the unique strategies employed by each strain to withstand exposure to very different concentrations of phosphine, but leading to equivalent rates of survival between the two strains.

Each of the first two analyses consisted of four treatments, only one of which was unique to each analysis – the *dld-1*(*wr4*) strain exposed to 500 ppm phosphine in the dose analysis and the *dld-1*(*wr4*) strain exposed to 4000 ppm phosphine in the “stress” analysis. Despite three of the data sets being common to both analyses (both strains in air and the N2 strain in 500 ppm phosphine), the results from the two analyses differed sharply. The number of genes whose expression levels changed due to exposure to phosphine decreased by 80% (5594 versus 1101). In contrast to the first analysis, in which 993 interacting genes were identified (i.e. genes that were differentially regulated according to both strain and exposure to phosphine), no such genes were each strain was exposed to the same level of phosphine-induced stress (i.e. their respective LC_50_s). This is a very strong indication that once the resistance mechanism is overcome, the response to an ultimately lethal dose of phosphine does not differ between the two strains. The implication is that resistance is achieved by an avoidance mechanism that reduces the toxicity of phosphine without fundamentally interfering with its mode of action.

A third analysis sought to identify constitutive differences in gene expression between the two strains. These genes might contribute to resistance mechanisms that do not rely on phosphine triggered changes in gene expression. For this analysis, a t-test was used to compare the expression levels of each gene between *N2* and *dld-1*(*wr4*), each of which had been cultured in the absence of phosphine. The expression levels of 256 genes differed between the two strains and passed the Benjamini and Hochberg multiple testing correction at a false discovery rate of 0.05 (Benjamini and Hochberg, 1995). 248 of these genes were selected for further analysis as their d expression levels differed by at least twofold.

We then analysed the 248 genes that are differentially expressed between the two strains in the absence of phosphine, together with the 993 genes that respond to phosphine differentially for their roles in cellular metabolism to allow us to determine how metabolism might influence the resistance phenotype. We employed several very different analytical approaches. Firstly, we developed a genome scale metabolic model that allowed us to visually inspect metabolic pathways for gene expression differences. The metabolic model had an added benefit as it allowed us to integrate gene expression differences with differences in the abundance of metabolites. Secondly, we used the more tradition gene set enrichment analysis to identify major processes associated with phosphine resistance. Thirdly, we simply inspected the data to see if they were consistent or inconsistent with proposed mechanisms of resistance gleaned from the literature (Nath et al. 2011).

### Construction of a *C. elegans* GSM for analysis and integration of the metabolite and gene expression data

We constructed a genome-scale model (GSM) of *C. elegans* metabolism, which we termed *Ce*Con (*<u>C. e</u>legans* re<u>Con</u>struction), to put the measured gene expression and metabolite level changes into context. *Ce*Con comprises 225 pathways, 1923 reactions and 1394 metabolites. It also identifies 20793 polypeptides, including 2754 enzymes and 73 transporters. Finally, the reconstruction contains 20791 protein coding genes. *Ce*Con relied on UniProt annotations which include localization of protein to different organelles, which provided our model with inbuilt compartmentalisation between cytosol, mitochondrion and lysosome. The following pathway-based analysis of gene expression and metabolite changes was obtained by using *Ce*Con as both an analytical and visualization tool.

### Survival of phosphine exposure by *dld-1*(*wr4*) is associated with global suppression of aerobic catabolism

#### Carbohydrate catabolism

In the transcriptomic and metabolomics data, we observed widespread suppression of aerobic catabolism in the *dld-1*(*wr4*) mutant strain exposed to 500 ppm phosphine, a sub-lethal dose for the mutant that would nevertheless kill the N2 strain. It was clear from the metabolic pathway analysis that genes encoding six enzymes of the TCA cycle, a central contributor to aerobic energy metabolism were down regulated. It is worth noting that **the *dld-1*(*wr4*) mutation causes partial loss of function in the enzyme dihydrolipoamide dehydrogenase (DLD)**, which is a subunit of **α-ketoglutarate dehydrogenase (KGDH)**, a multi-enzyme complex that is also the rate limiting step of the TCA cycle. The six TCA cycle genes that were down regulated included *ogdh-1,* the gene encoding the E1 subunit of the KGDH complex. In addition, the levels of two metabolites of the TCA cycle, fumarate and succinate, decrease strongly.

The glyoxylate cycle, which provides a means of bypassing KGDH, is found in bacteria, fungi, plants and nematodes. The gene for the defining enzyme, *icl-1* (actually a dual function isocitrate lyase-malate synthase enzyme in *C. elegans*), is up regulated in response to 500 ppm phosphine, particularly in N2, but also in the mutant. This could be a strategy to bypass KGDH to compensate for the metabolic inhibition caused by phosphine exposure. One of the products of the isocitrate lyase-malate synthase enzyme is succinate. The level of the succinate metabolite is very low in both strains in response to phosphine exposure, but it is indeed higher in the N2 strain.

Another metabolic pathway that bypasses KGDH is the GABA shunt. Rather than glutamate being converted to α-ketoglutarate and ultimately to succinate via KGDH, the GABA shunt converts glutamate to succinate via gamma amino butyric acid (GABA). The levels of glutamate increase in wild type *C. elegans* in response to either sublethal or toxic levels of phosphine. In contrast, the same effective exposure top phosphine causes the glutamate levels to drop in the *dld-1* mutant strain. The gene expression data provides no evidence of genes of the GABA shunt being differentially regulated by strain or by exposure to phosphine. The source of the observed differences in glutamate levels remains to be discovered, but there are a variety of pathways that act on glutamate other than the GABA shunt.

**Figure Legend:**
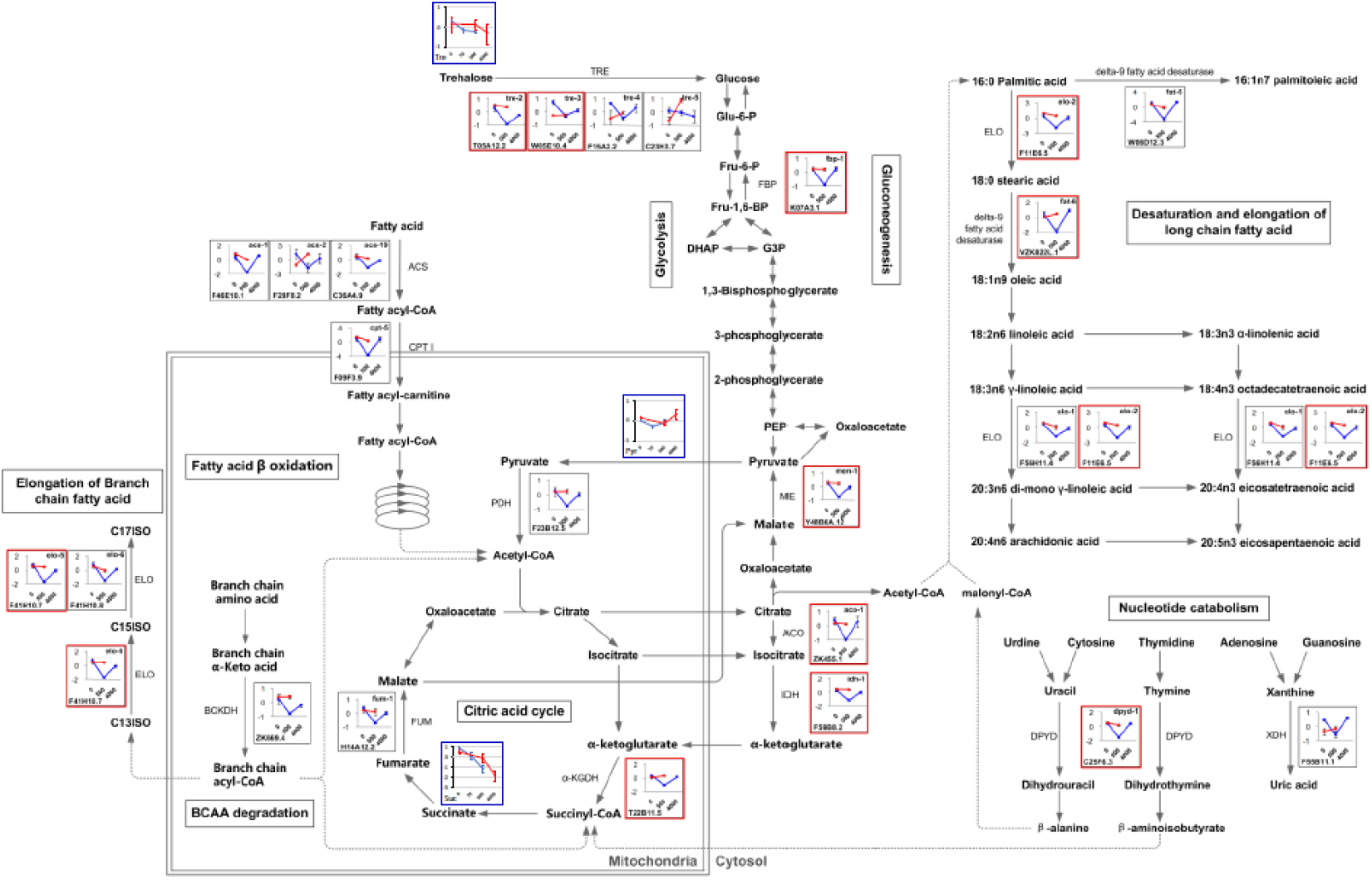
6 steps in the TCA cycle and 3 steps in the Glyoxylate cycle exhibited decrease in excess of a two-fold difference.

It is interesting to note that there is no indication that glycolysis is inhibited as none of the glycolytic genes showed any appreciable change in expression in either N2 or the *dld-1*(*wr4*) mutant strain. Importantly, phosphine did not affect expression of either of the two phosphofructokinase genes. As phosphofructokinase catalyzes the committed step of glycolysis, this supports the notion that anaerobic energy metabolism is not inhibited.

We do find that the *dlat-1* gene is suppressed specifically in the *dld-1*(*wr4*) mutant when exposed to 500 ppm phosphine. This gene encodes the E2 subunit of the **pyruvate dehydrogenase complex (PDH)**, the enzyme complex that feeds the pyruvate end product of glycolysis into the TCA cycle for aerobic respiration. Like KGDH, the PDH complex has DLD as one of its subunits. Thus, it appears that, whereas glycolytic gene expression is not affected by either the *dld-1*(*wr4*) mutation or exposure to phosphine, the activity of PDH is likely to be strongly impacted both by the *dld-1*(*wr4*) mutation and by suppression of *dlat-1* gene expression caused by exposure to 500 ppm phosphine. As a result, the delivery of acetyl CoA to the TCA cycle via glycolytically derived pyruvate is almost certainly impaired in the mutant, particularly in the presence of 500 ppm phosphine. The pyruvate level increased modestly during phosphine exposure in the N2 strain, but actually decreased in the *dld-1*(*wr4*) mutant. This difference between the two strains and deviation from the expected response for the mutant strain indicates that the mutant nematodes have adapted to the dld-1 mutation in a way that protects the PDH complex from the influence of phosphine or that shunts the pyruvate that cannot be catabolized through PDH to another catabolic pathway.

There are several alternative fates of pyruvate. PDH inhibition in humans results in lactic acidosis, but we see that phosphine exposure results in a decrease in lactate levels which is especially pronounced in the *dld-1*(*wr4*) mutant. Gluconeogenesis is a sink for pyruvate, but key genes in that pathway are suppressed as described below. Alanine levels do increase suggesting that a pyruvate transaminase could be activated, but this is not apparent according to transcript abundance.

The pentose phosphate pathway provides an alternative mechanism for the complete catabolism of glucose to CO_2_ that bypasses both glycolysis and the TCA cycle with the generation of NADPH rather than NADH. There is no evidence that the pentose phosphate pathway genes are either upregulated to compensate for the down regulation of the TCA cycle or downregulated in concert with the down regulation of the TCA cycle.

#### Amino acid catabolism

Heterogeneous changes in the levels of arginine, aspartate, asparagine, histidine, lysine, phenylalanine, proline, threonine, tryptophan, and tyrosine were observed in the metabolomic data. Tryptophan levels showed a behavior reminiscent of the branched-chain amino acids (BCAAs), which has been observed in other biological processes, where levels of tryptophan and/or phenylalanine cluster with BCAAs (Mansfeld et al., Newgard et al., Fuchs et al. 2010)

We also find that the *dbt-1* gene is suppressed specifically in the *dld-1*(*wr4*) mutant when exposed to 500 ppm phosphine. This gene encodes the E2 subunit of the **branched chain ketoacid dehydrogenase complex (BCKDH)**, a DLD containing enzyme complex that participates in the catabolism of the BCAAs leucine, isoleucine and valine, so suppression of the *dbt-1* gene should disrupt normal catabolism of these amino acids. The levels of the three BCAAs exhibit though a different behavior than pyruvate and glutamate: In the untreated mutant they are consistently lower than in the wild-type (reflecting constitutive downregulation of this pathway), then at sub-lethal phosphine concentrations increase sharply in the mutant while staying level or decreasing in the wild-type, where they only increase at the LC_50_ (500 ppm).

We also find that the *gcsh-2* gene is suppressed specifically in the *dld-1*(*wr4*) mutant when exposed to 500 ppm phosphine. This gene encodes the H subunit of the **glycine cleavage system (GCS)** a DLD containing enzyme complex that participates in the catabolism of glycine (Figure 3). Glycine levels increase in both wt and mutant nematodes at sub-lethal phosphine concentrations. At LC_50_ phosphine levels, glycine levels stay high in the wild-type while decreasing in the mutant to below baseline levels. This runs counter to the gene expression results, but alternative pathways exist for glycine catabolism, including a paralogue of *gcsh-2* that was not present in the microarray.

#### Fatty acid catabolism

Response of the phosphine resistant mutant *dld-1* to a dose of phosphine (500 ppm) equivalent to the LC50 of the parental wild type strain *N2*, results in a coordinated transcriptional suppression of catabolic pathways. Beta-oxidation of fatty acids in the mitochondria is the most strongly inhibited of these pathways. To pass through the mitochondria inner membrane, fatty acids must first be activated by the addition of acyl-CoA via acyl-CoA synthetase in the cytosol. Activated fatty acids are then converted to fatty acyl-carnitine by the carnitine palmitoyl transferase. The fatty acyl-carnitine then can be transported across the inner mitochondrial membrane into the matrix. Conversion to the carnitine ester commits the fatty acid to the oxidative fate. The carnitine-mediated entry process is the rate-limiting step for oxidation of fatty acids in mitochondria. The activation of free fatty acids by addition of coenzyme A, is also inhibited strongly as seen in the significant suppression of three different isoforms of acyl-CoA synthetase, with acs-2 inhibited approximately 7 fold. One carnitine-palmitoyl transferase 1 isoform (cpt-5) is suppressed to a much greater extent, ~25 fold. By targeting the most rate limiting steps, only a small subset of genes need to be regulated to control the activity of the entire pathway.

#### Nucleotide catabolism

As with the other biosynthetic pathways, nucleotide catabolism is also suppressed in *dld-1* when exposed to 500 ppm phosphine (Figure 2). Specifically, dihydropyrimidine dehydrogenase, the rate limiting step of pyrimidine catabolism is inhibited in *dld-1* at the sublethal concentration of 500 ppm phosphine, but it is not inhibited when either *N2* or *dld-1* are exposed to phosphine at their LC50s. A similar situation exists for purine catabolism, as the gene encoding the rate limiting enzyme, xanthine dehydrogenase is also suppressed in *dld-1* at 500 ppm. In this case, *N2* expression is constitutively low, whereas expression of the gene increases at the LC50 of *dld-1.*

**Figure 2:**
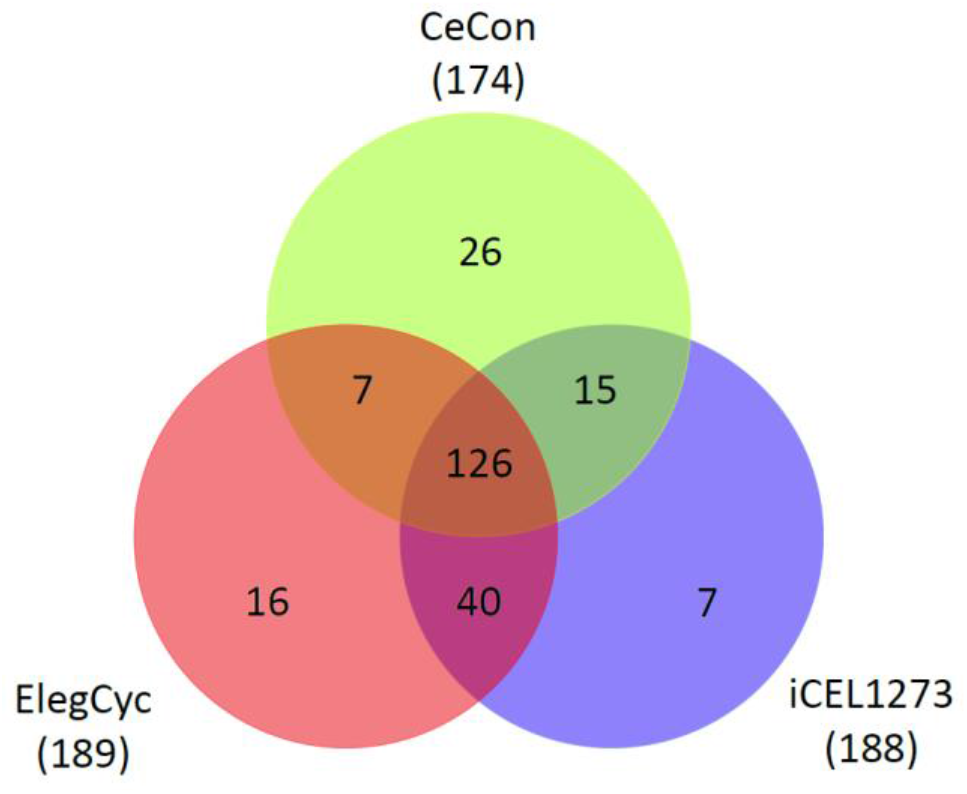
Pathway comparisons between CeCon, ElegCyc and iCEL1273.

### Survival of phosphine exposure by *dld-1* is associated with global suppression of major biosynthetic pathways

#### Fatty acid synthesis and gluconeogenesis

Suppression of core biosynthetic pathways in *dld-1* occurs in parallel with the suppression of catabolism when the nematodes are exposed to 500 ppm phosphine (Figure 2). This includes suppression of elongation of branched chain fatty acids and long chain fatty acids as well as suppression of fatty acid desaturation. It also includes suppression of genes for the metabolism of TCA cycle intermediates that have been transported from the mitochondrial matrix to the cytoplasm.

These pathways are responsible for the cytoplasmic synthetic pathways. For instance, the acetyl-CoA generated by these reactions contributes to fatty acid biosynthesis. As a result, suppression of these genes likely contributes to the suppression of fatty acid biosynthesis. Suppression of these pathways can also limit the availability of pyruvate for gluconeogenesis. Likewise, the gene that encodes fructose-1,6-bisphosphatase, the rate limiting step for gluconeogenesis, is also suppressed. Thus it seems that synthesis of the major energy metabolites, i.e. fats and sugars, is likely to be suppressed.

### Oxidative Stress

Oxidative stress due to mitochondria-generated reactive oxygen species (ROS) is the prevailing model of phosphine toxicity. We examined the gene expression data to determine whether the phosphine resistance of *dld-1* is due to the induction of an oxidative stress response system. Specifically, we looked for induction of superoxide dismutase, catalase and peroxidase genes as these genes are able to detoxify the primary reactive oxygen products of mitochondrial respiration. If defence against the toxic effects of ROS is indeed a mechanism of resistant toward phosphine, then antioxidant genes should be more strongly induced in *dld-1* than in *N2.* If the *dld-1* strain simply suffers less oxidative stress than *N2* strain during phosphine exposure, as might happen if uptake is inhibited, these antioxidant genes should be more strongly induced in *N2* than in *dld-1.*

Five genes encode superoxide dismutase in *C. elegans,* two of which, sod-1 and sod-2 show a significant strain x dose interaction. The magnitude of the change in gene expression between two strains is much less than 2 fold, however. Only two sod genes, sod-4 and sod-5, are induced more than 2 fold by phosphine, but the induction is not significantly different between the two strains. sod-4 is an extracellular isoform proposed to be involved in the signalling activity of ROS, whereas sod-5 is cytoplasmic. These genes are induced approximately 3 fold when each strain is exposed at their LC50, but less than 2 fold when *dld-1* is exposed to phosphine at the sublethal concentration of 500 ppm. The lower level of expression at the sublethal dose clearly indicates that induction of sod genes is not responsible for the ability of the resistant mutant to survive at 500 ppm.

Among the 3 genes that encode catalase, none showed a significant strain x dose interaction at 500 ppm. Furthermore, the magnitude of inductions between two strains differed by less than 2 fold. Among the 3 peroxidases, prdx-6 showed an interaction effect, but the magnitude of inductions between the two strains differed by less than 2 fold. Thus, it seems that none of the genes that might detoxify superoxide or hydrogen peroxide were differentially regulated to an extent that would explain the phosphine resistance characteristics of the two strains. Furthermore, the basal expression level of none of these genes differed by more than 2 fold between the two strains. This implies that the phosphine resistance of *dld-1* is not due to a constitutively inducted antioxidant defense system in this strain.

Gamma-glutamine cysteine synthetase (gcs-1) catalyzes the rate-limiting step in glutathione biosynthesis. Under same dose 500 ppm exposure, gcs-1 showed no interaction effect between strain and dose.

### Metabolism of sulfur-containing amino acids

Metabolism of the two sulfur-containing, proteogenic amino acids, methionine and cysteine, impacts several critical biological systems. Methionine is key to single carbon metabolism as it contributes directly to the biosynthesis of two different methyl donors, 5,10-methylene tetrahydrofolate (MTHF) and S-adenosylmethionine (SAM). Methionine also contributes to sulfur metabolism through SAM and the biosynthesis of cysteine. Cysteine contributes to redox homeostasis through glutathione, which is the primary cellular redox regulator. Cysteine metabolism can also give rise to the gasotransmitters hydrogen sulfide (H_2_S) and sulfur dioxide as well (Li et al., 2009).

Overall, it is clear that exposure of the resistant mutant to 500 ppm phosphine results in suppression of enzymes throughout all aspects of single carbon metabolism and sulfur metabolism as well (Figure 3). In contrast, exposure of either wild type nematodes or the resistant mutant to phosphine at their respective LC_50_ values, results in gene expression across these pathways that is the same as in control nematodes that have not been exposed to phosphine.

**Figure 3:**
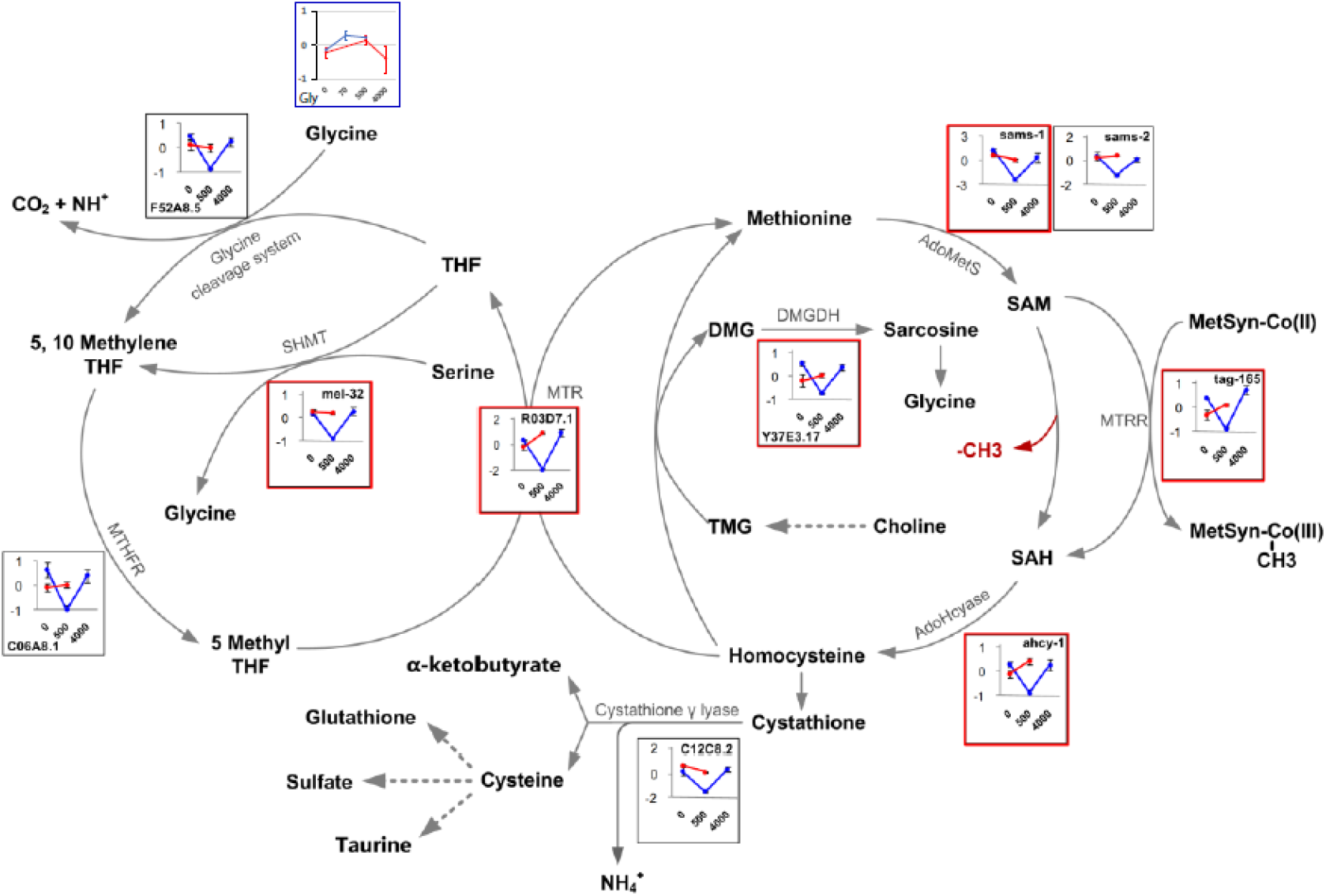
Gene expression and metabolite changes connected to the glycine cleavage system and one-carbon metabolism.

Many of the enzymatic steps are represented by multiple isoforms, but usually only a single gene is suppressed in the resistant mutant at 500 ppm. This is sometimes due to the stringent criteria that we have set for inclusion in the analysis, i.e. an interaction p-value of <0.05 (corrected for multiple testing) as well as a 2-fold difference in expression change between the two strains. In other cases, however, paralogous genes are differentially regulated, possibly related to distinct functions or differential patterns of subcellular, cellular or tissue distribution. In this regard, it is interesting that cystathionine gamma lyase (CSE) is suppressed more strongly than cystathionine beta synthase (CBS) (suppressed in the mutant by exposure to 500 ppm phosphine. Both of these enzymes can produce the gasotransmitter H2S, but CSE does this in liver and smooth muscle in mammals, whereas CBS generates H2S in the brain. An enzyme for mitochondria specific H2S synthesis, MTS, is not suppressed in the mutant at low dose. Cystathionine levels themselves are almost unaffected by phosphine exposure in wild-type *C. elegans,* while in the mutant they increase at sublethal phosphine dose and drop below baseline at the LC_50_.

Despite the fact that there is some uncertainty regarding the tissue specificity, developmental timing or subcellular localisation of gene regulation, it is quite clear that suppression of methionine and cysteine metabolism as well as associated metabolic pathways occurs quite specifically when the resistant mutant is exposed to a sublethal dose of phosphine. Thus, it seems likely that methyl transfer, sulfur transfer, the cellular redox state and polyamine biosynthesis is all significantly affected by phosphine in the resistant mutant.

### Xenobiotic response genes

After a xenobiotic compound gets entered into the cell, organisms have the enzymatic and transporting system to excrete the compound out of the cell so as to avoid accumulation. Most of the xenobiotic-metabolizing enzymes are localized in the endoplasmic reticulum, known as phase I enzymes such as cytochrome P450 (CYP), flavin-containing monooxygenase (FMO), and also phase II enzymes like UDP-glucuronosyltransferase (UGT) and glutathione S-transferase (GST) (Cribb et al. 2005). After biotransformation by these two major pathways, the compound is then actively transferred out of the cell through efflux systems in the cell membrane such as ATP-binding cassette (ABC) and major facilitator superfamily (MFS) transporters (Hayashi 2003). It is also worth to note that in many cases biotransformation conducted by phase I and phase II enzymes can lead to the formation of more toxic intermediates than the original compound (Cribb et al. 2005).

Microarray data indicated same dose of phosphine exposure 500 ppm differentially regulated a number of family members of these enzymes between the two strains:

1. 10 phase I enzymes, 13 phase II enzymes and 10 phase III transporters showed more dramatic induction in dld-1(wr4) than that in N2.
2. 2 phase I enzymes, 10 phase II enzymes and 14 phase III transporters showed more dramatic suppression in dld-1(wr4) than that in N2.
3. 7 phase I enzymes, 11 phase II enzymes and 10 phase III transporters showed more dramatic induction in N2 than that in dld-1(wr4).
4. There are no genes showed more dramatic suppression in N2 than that in dld-1(wr4).

Overall, since the induction of these xenobiotic metabolizing genes were not particularly enriched in one of the strains, thus before we get to know the individual function and the tissue specificity of each of these enzymes, at this stage it is difficult to draw any conclusion on how these xenobiotic metabolizing genes contribute to dld-1(wr4)’s resistance or N2’s sensitivity under the same dose exposure 500 ppm. In addition, the suppression of several xenobiotic metabolizing genes observed in dld-1(wr4) might be associated with other functions of these enzymes, such as endogenous hormone production.

## Discussion

We have characterised the response of wild-type and resistant C. elegans with transcriptomic and metbaolomic methods. In addition, we have constructed a GSM of *C. elegans* that serves as a tool to interpret the experimental data. Genome-scale modeling (Palsson 2006), beyond enabling the prediction of metabolic fluxes allows to derive context-specific metabolic networks from –omics datasets (transcriptomics, metabolomics, proteomics) that reflect metabolic activity on an organism-wide scale for conditions of interest (Gebauer et al. 2016). Based on such networks, key enzymes responsible for metabolic adaptation as well as interventions to counteract them can be identified (Yizhak et al. 2013). Thus, using CeCon to analyze data provides visual output in the context of the metabolic pathways of *C. elegans* which aids in recognition and interpretation of the biology. Some connections and patterns become more readily apparent that would otherwise be more difficult to identify using other means of analysis.

Indeed, analyzing the phosphine response in the context of the GSM can shed light on which potential resistance mechanisms are possible and which are unlikely. Originally the following mechanisms were considered as contenders for explaining phosphine resistance in *C. elegans*:

1. Mitochondrial inhibition that results in the generation of reactive oxygen species (ROS)
2. Hypometabolism, i.e. protective metabolic suppression and a decrease in metabolic rate.
3. Using the xenobiotic response system to counteract phosphine toxicity

Analysis of the gene expression data found no significant alteration of ROS response genes, and data were inconclusive on an involvement of the xenobiotic response system. The most striking feature of the phosphine-resistant *C. elegans* dld-1(wr4) strain is though the constitutive downregulation of its central core metabolism, but especially of the TCA cycle and associated pathways. This state of hypometabolism is the most likely mechanistic explanation for phosphine resistance.

Indeed, oxidative respiration in *the dld-1*(*wr4*) strain is constitutively suppressed to about 25% of the normal rate of respiration (Zuryn et al., 2008). Phosphine exposure causes a further decrease in the rate of oxygen consumption in *dld-1*(*wr4*) from 25% to about 15% of normal (Zuryn et al., 2008), which is likely to be associated with the changes that we observe in the expression of catabolic genes. In contrast, the wild-type *N2* strain responds quite differently, as phosphine exposure does not alter the expression of genes involved in aerobic energy metabolism. Despite the lack of change in gene expression, aerobic respiration drops to about 15% of normal levels, most of which occurs within 60 minutes of exposure to phosphine. The speed and the magnitude of the respiratory inhibition indicate that metabolism is suppressed via a non-transcriptional mechanism, e.g. normal regulation of enzyme activity through metabolite levels (see below). (Insert Ma Li’s reference here?) The low oxygen consumption is likely to be the consequence of this metabolic restructuring into hypometabolism. It decreases the level of toxic stress that results from phosphine exposure, and might thus provide the opportunity for the transcriptionally orchestrated metabolic restructuring observed in *dld-1*(*wr4*), but not *N2.*

The *dld-1*(*wr4*) strain of *C. elegans* is already in a hypometabolic state and preadapted to phosphine exposure. The only difference between this strain and the wild type is a point mutation in the gene for DLD. That means that DLD must be responsible for the hypometabolic state of the mutant. Indeed, DLD is suited to this role: It is present at four key points of the metabolic network. That means any alteration of the activity or function of DLD has wide-ranging consequences for metabolism. Intriguingly, DLD is known to be competitively product inhibited by NADH (Sahlman & Williams 1989, Wilkinson & Williams, 1981). That means that DLD is effectively a sensor for the mitochondrial redox potential. Furthermore, the E2 subunit of PDH is product inhibited by AcCoA, making it sensitive to energy flux. Thus, any change in both parameters will immediately feed back into the metabolic network, making DLD and its associated enzyme complexes potential master regulators of metabolic activity.

In parallel with *Ce*Con, three other *C. elegans* GSMs were constructed independently. Two of them have been published, ElegCyc and iCEL1273 (Gebauer et al. 2016, Yilmaz & Walhout 2016). Additionally, various public but unpublished metabolic reconstructions exist for C. *elegans* such as SolCyc (*http://solcyc.solgenomics.net*) and BMID000000141468 (Buchel et al. 2013). The development of multiple models for a single organism is common, an example being Saccharomyces cerevisiae, with over two dozen metabolic models (Yilmaz & Walhout 2017). As models of the same organism are generated to study a particular characteristic or application, they complement each other, thus providing a more complete overview of the information available (van Heck et al. 2016). For example, the human metabolic models share only a 3% consensus in their reactions (Stobbe et al. 2011). Indeed, a comparison of CeCon with the other models shows a similar situation:

An initial comparison of the coverage and size of the metabolic models (Table 1) shows that ElegCyc contains the largest number of pathways and transcription units, whereas CeCon contains many more reactions and metabolites, as well as gene relations.

**Table 1:** A top-level comparison of model size between different *C. elegans* GSMs.

| Model | *CeCon* | *ElegCyc* | *iCEL1273* | *SolCyc* | **BMID00000141468** |
| --- | --- | --- | --- | --- | --- |
| **Pathways** | 225 | 273 | 100 | 166 | 0 |
| **Enzymatic Reactions** | 1907 | 1171 | ~ | 1085 | 1119 |
| **Transport Reactions** | 16 | 22 | 475 | 13 | 1162 |
| **Combined Reactions** | ~ | ~ | 1985 | ~ | ~ |
| **Polypeptides** | 20793 | 24129 | ~ | 20433 | 17 |
| **Protein Complexes** | 0 | 66 | ~ | 1 | 0 |
| **Enzymes** | 2754 | 1694 | 623 | 1779 | 937 |
| **Transporters** | 73 | 27 | ~ | 43 | 0 |
| **Compounds** | 1394 | 773 | 887 | 755 | 996 |
| **Genes** | 20791 | 4620 | 1273 | 20430 | 0 |
| **Protein Coding Genes** | 20791 | 4419 | ~ | 20430 | 0 |
| **Transcription Units** | ~ | 21213 | ~ | 0 | 0 |
| tRNAs | ~ | 696 | ~ | 0 | 0 |
| Compartments | 3 | 5 | 3 | ~ | 3 |

A more fine-grained comparison between *Ce*Con and the two published GSMs revealed some consensus. 237 pathways were identified across all three models, with duplicate pathways grouped under the same name. ElegCyc and iCEL1273 share 70% of these 237 pathways in the analysis. Comparing CeCon to ElegCyc and iCEL1273, pathway similarity is 56% and 59% respectively. 49 of pathways in the analysis (21% of 237 pathways) were found to be unique to a single model, with CeCon containing the greatest number of unique pathways.

**Table 2:**
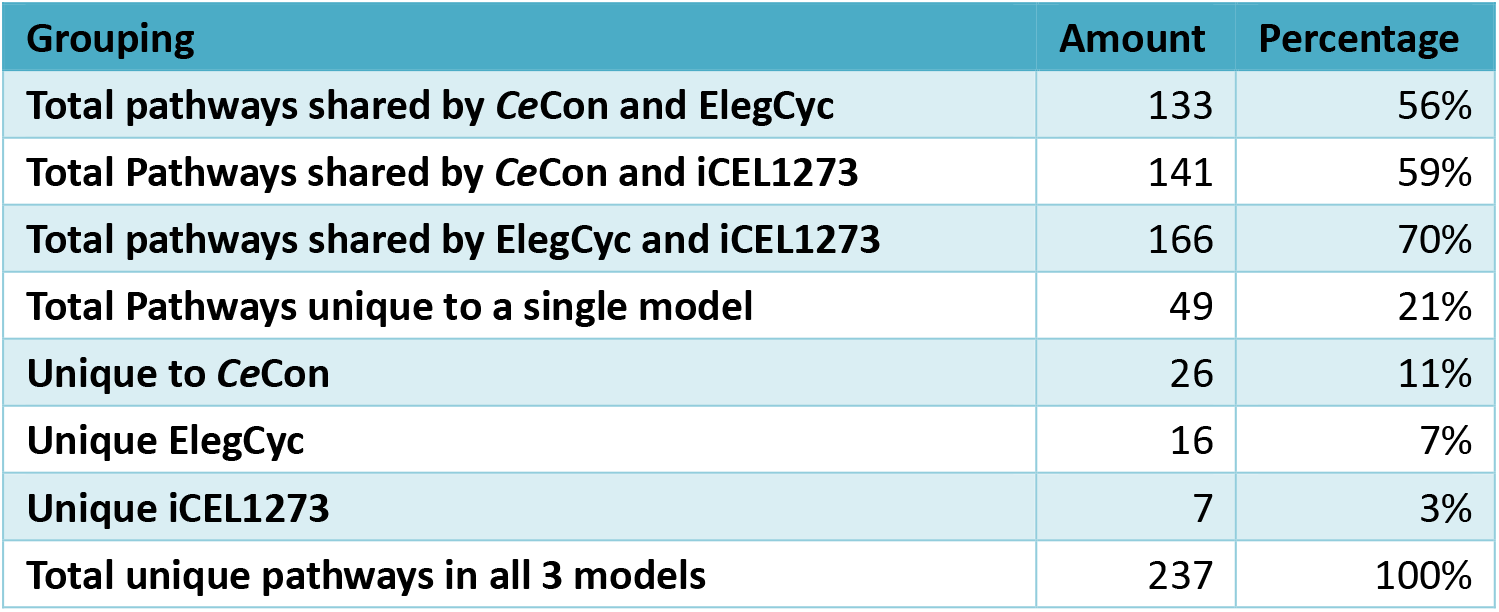
Pathway comparisons between *Ce*Con, ElegCyc and iCEL1273.

| Grouping | Amount | Percentage |
| --- | --- | --- |
| Total pathways shared by CeCon and ElegCyc | 133 | 56% |
| Total Pathways shared by CeCon and iCEL1273 | 141 | 59% |
| Total pathways shared by ElegCyc and iCEL1273 | 166 | 70% |
| Total Pathways unique to a single model | 49 | 21% |
| Unique to CeCon | 26 | 11% |
| Unique ElegCyc | 16 | 7% |
| Unique iCEL1273 | 7 | 3% |
| Total unique pathways in all 3 models | 237 | 100% |

26 (11%) unique pathways were found in our model, with ElegCyc and iCEL1273 containing 16 (7%) and 7 (3%) respectively. This greater individuality is likely due to the protein-based approach used to generate the model. The pathway comparison can be found in the supplementary information [Supplementary File.xlsx].

At a reaction granularity, the MetaNetX namespace comparison revealed that only 261 (7%) reactions from CeCon exactly matched a reaction in ElegCyc and 239 (6%) of reactions matched in iCEL1273. Similarly, only 6% reactions were matched exactly in metabolites, stoichiometry and directionality between ElegCyc and iCEL1273.

The fact that there are now three actively curated *C. elegans* GSMs, which overlap only partially, argues strongly for the future generation of a consensus GSM of *C. elegans,* similar to the work carried out on the human consensus GSM. An initial workshop to commence this process was held in April 2017 in Cambridge (UK), foreshadowing rapid developments in the future in this area (Hastings et al. 2017).

## Conclusions

In this study we investigated the mechanism of phosphine resistance in *C. elegans* using transcriptomics and metabolomics. In addition, we constructed *Ce*Con, a C. elegans GSM, as a useful tool for analyzing the experimental –omics data and putting them into their metabolic context. Together with the other available C. elegans GSMs, CeCon also forms the basis for a future consensus GSM of the nematode.

Our analysis shows that hypometabolic adaptation is the main phosphine resistance mechanism. As the mutation in DLD is the only change between the wild-type and the resistant *dld-1* strain, it means DLD is implicated in orchestrating this hypometabolic adaptation. This implies a potential role for DLD in other phenomena that involve hypometabolism such as reperfusion injury and metabolic resistance.

## List of abbreviations

ANOVA: Analysis of variance
BCAA: branched-chain amino acid
BCKDH: branched-chain ketoacid dehydrogenase
*Ce*Con: *<u>C. e</u>legans* re<u>con</u>struction
CPMG: Carr-Purcell-Meiboom-Gill
CSE: cystathionine gamma lyase
DLD: dihydrolipoamide dehydrogenase
DSS: 2,2-dimethylsilapentane-5-sulfonic acid
GCS: glycine cleavage system
GABA: gamma amino butyric acid
GO: gene ontology
GSM: genome-scale metabolic model
HMBC: heteronuclear multiple bond correlation
HSQC: heteronuclear single quantum correlation
IMB: Institute for Molecular Bioscience
KGDH: alpha-ketoglutarate dehydrogenase
LC_50_: Medial lethal concentration
MTC: multiple testing correction
MTHF: 5,10-methylene tetrahydrofolate
NMR: nuclear magnetic resonance
NOESY: nuclear Overhauser effect spectroscopy
PDH: pyruvate dehydrogenase
ROS: reactive oxygen species
SAM: S-adenosyl methionine
TCA: tricarboxylic acid
TOCSY: total correlation spectroscopy
wt: wild-type

## Declarations

### Ethics approval and consent to participate

No ethics approvals were needed for *C. elegans* work.

### Consent for publication

Not applicable

### Availability of data and material

All data analysed or generated during this study are available from the authors on reasonable request. Raw NMR metabolomics data were deposited in the *MetaboLights* database (*http://www.ebi.ac.uk/metabolights/*). The *C. elegans* GSM CeCon is accessible online at *http://cai-gsmodel.cai.uq.edu.au*.

## Funding

This study was supported by funding from the Plant Biosecurity Collaborative Research Centre (PBCRC), established between the Australian Research Council and the Australian grains industry. PRE received project funding from the PBCRC and the research appointment of HJS was partially funded by the PBCRC.

The PBCRC had no role in the design of the study; in collection, analysis, and interpretation of data; and in writing the manuscript. The final manuscript version was submitted to the PBCRC as milestone deliverable.

## Authors’ contributions

LM: Designed, carried out, analysed the microarray experiments, and contributed to draft of the manuscript.

AHCC: Constructed *Ce*Con, and contributed to the draft of the manuscript.

JH: Refined CeCon and compared it with other *C. elegans* GSMs, contributed to the manuscript draft and to several figures.

PRE: Conceived the study, designed the major experiments, contributed to the biological interpretation, drafted and refined the manuscript.

HJS: Designed, carried out, and analysed the NMR-based metabolomics experiments, contributed to the biological interpretation, contributed to figures, drafted and refined the manuscript.

All authors read and approved the final manuscript.

## Acknowledgements

We thank Mr Nicholas Mok for help in harvesting and extracting worms for the metabolomics experiments.

## Supplementary

### Metabolomics Samples

**Supplementary Table 1:**
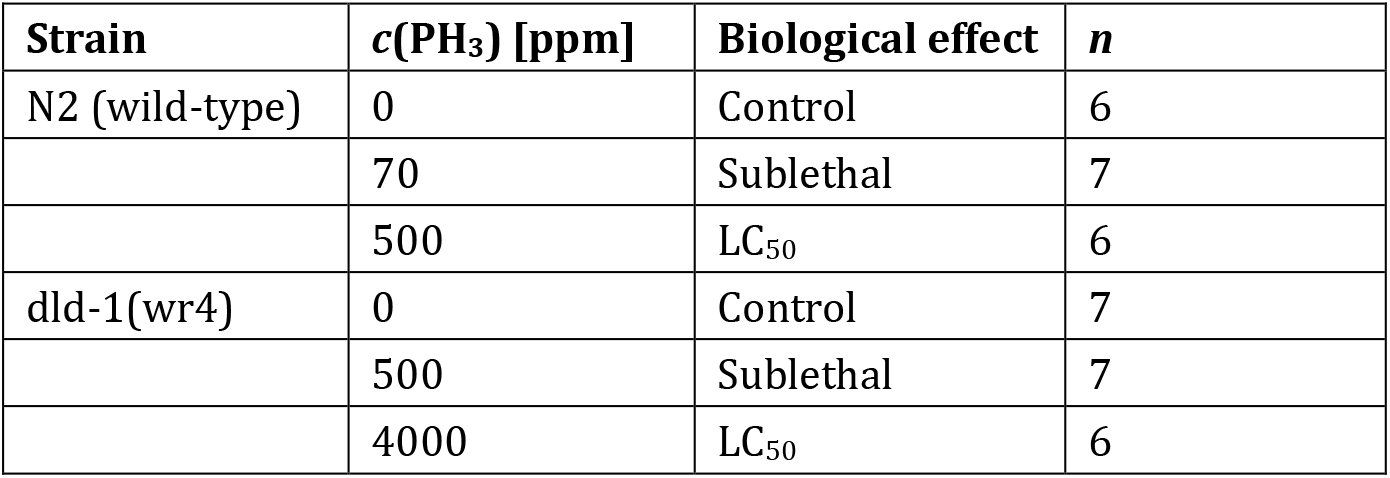
*C. elegans* samples used in the metabolomics experiments. *c*(PH_3_)= phosphine concentration in ppm, *n*= number of samples.

| Strain | $c(\text{PH}_3)$ [ppm] | Biological effect | $n$ |
| --- | --- | --- | --- |
| N2 (wild-type) | 0 | Control | 6 |
|  | 70 | Sublethal | 7 |
|  | 500 | LC<sub>50</sub> | 6 |
| dld-1(wr4) | 0 | Control | 7 |
|  | 500 | Sublethal | 7 |
|  | 4000 | LC<sub>50</sub> | 6 |

### Benchmarking

To benchmark the effectiveness of our database generation methods, we generated a metabolic map for human metabolism using UniProt data and compared it against the latest manually curated version, the summation of over ten years worth of work. Coverage indicators such as the number of pathways, enzymatic reactions, enzymes, and compounds involved were compared between the two databases.

The coverage of our database exceeded the coverage of the manually curated one by 101% to 691%, depending on the metric used.

**Supplementary Table 2:** A comparison of the coverage of the metabolic databases generated through manual and automatic curation respectively. Our method was able to produce a database with a 102% to 691% coverage rate of its manually curated counterpart.

| Metric | Manual | Automatic | Comparison |
| --- | --- | --- | --- |
| Pathways | 289 | 373 | 129.07% |
| Enzymatic Reactions | 2324 | 2840 | 122.20% |
| Enzymes | 393 | 2712 | 690.08% |
| Compounds | 1798 | 1832 | 101.89% |

### Annotation Sources

During the development of our database, we primarily utilized data from the UniProt database. Coverage of the metabolic map was increased by 12% to 46% (depending on the metric used) when the UniProt data was supplemented with annotations from the WormBase database, primarily GO terms.

A proteome by proteome BLAST was then conducted to compare *C. elegans* proteins against those of other major model organisms, namely H. sapiens, D. melanogaster, S. cerevisiae S288C and E. coli K12. In order to normalize against the effects of differing proteome sizes, the best matches were then BLAST-searched against the *C. elegans* proteome and the reciprocal E-value of the original BLAST searches was used as an indicator for which organism was the closest.

*C. elegans* genes for which there was missing information and a strong match in other organisms was further supplemented with data from that organism. 719 EC numbers were borrowed from S. cerevisiae matches. With this additional supplementary information, improvement levels reached 21% to 58% of the original.

**Supplementary Table 3:**
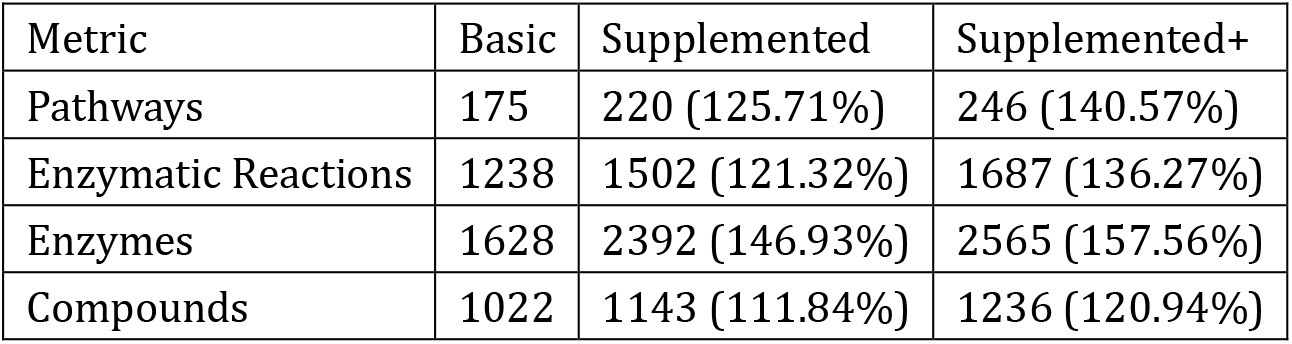
A comparison of the coverage of the *C. elegans* databases generated using different levels of annotation depth. The most basic metabolic map (Basic) was generated using only *C. elegans* data from the UniProt database. The second level map (Supplemented) was generated using the basic metabolic map, supplemented with Wormbase annotations. The third level map (Supplemented+) consisted of the data used to generate the second level map, further supplemented with annotations from BLAST matches for *C. elegans* genes that were deficient in annotations.

**Supplementary table 4:**
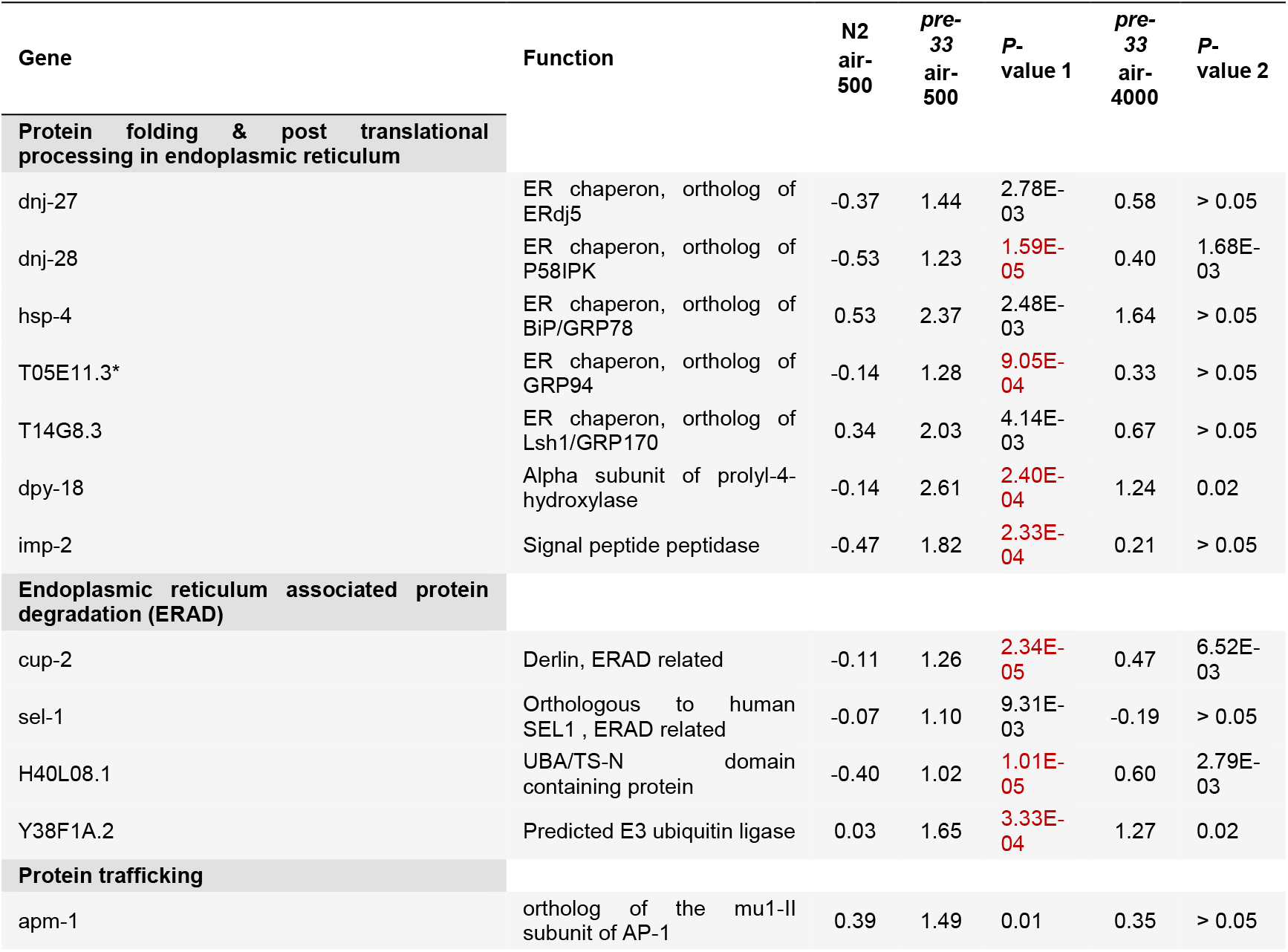
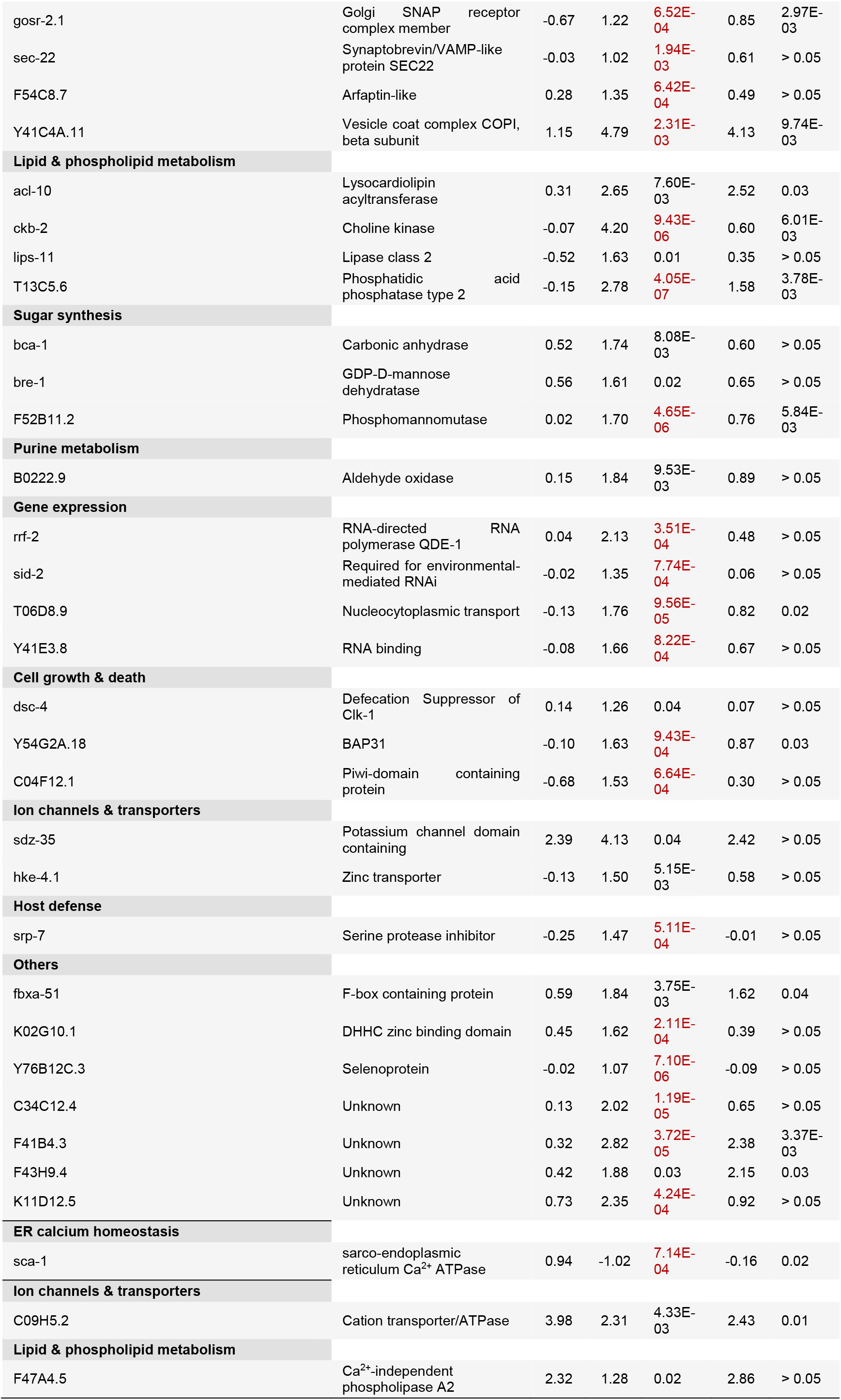

**Supplementary table 5:**

| Gene | Function | N2<br>air-500 | pre-33<br>air-500 | p-value 1 | pre-33<br>air-4000 | p-value 2 |
| --- | --- | --- | --- | --- | --- | --- |
| ins-2 ($\beta$) | Unknown | -0.18 | 1.02 | 1.01E-03 | -0.11 | > 0.05 |
| ins-12 ($\gamma$) | Unknown | 0.70 | 2.90 | 5.19E-04 | 1.30 | > 0.05 |
| ins-21 ($\alpha$) | Unknown | -0.59 | 1.14 | 8.38E-03 | -0.27 | > 0.05 |
| ins-9 ($\beta$) | Unknown | 0.13 | -1.56 | 0.03 | -1.36 | > 0.05 |
| ins-33 ($\alpha$) | Unknown | -1.18 | -3.82 | 1.59E-03 | -2.06 | > 0.05 |
| ins-39 ($\alpha$) | DAF-2 agonist | -0.73 | -3.16 | 4.54E-04 | 0.01 | > 0.05 |
| daf-28 ($\alpha$) | DAF-2 agonist | -0.57 | -1.60 | 2.52E-03 | -0.29 | > 0.05 |

**Supplementary table 6:**

| Gene | N2<br>air-500 | pre-33<br>air-500 | P-value 1 | pre-33<br>air-4000 | P-value 2 |
| --- | --- | --- | --- | --- | --- |
| nhr-38 | -1.40 | 2.35 | 3.21E-04 | 0.37 | 0.02 |
| nhr-54 | 0.60 | 1.73 | 0.05 | 0.85 | > 0.05 |
| nhr-67 | -0.66 | 1.91 | 3.41E-03 | 0.66 | > 0.05 |
| nhr-111 | -0.12 | 1.25 | 6.70E-03 | 0.65 | > 0.05 |
| nhr-147 | 0.01 | 1.79 | 9.81E-03 | 0.38 | > 0.05 |
| nhr-186 | -0.68 | 1.72 | 3.95E-04 | 0.08 | > 0.05 |
| nhr-241 | -0.42 | 1.09 | 7.94E-04 | 0.13 | > 0.05 |
| ZK418.10 | -0.69 | 1.01 | 9.45E-03 | -0.55 | > 0.05 |
| nhr-1 | -0.17 | -1.50 | 5.75E-04 | -0.30 | > 0.05 |
| nhr-31 | 0.37 | -1.55 | 8.39E-05 | -0.07 | > 0.05 |
| nhr-55 | 0.47 | -1.38 | 1.86E-04 | 0.61 | > 0.05 |
| nhr-114 | -0.12 | -1.64 | 0.01 | 0.59 | > 0.05 |
| nhr-143 | 0.63 | -1.10 | 0.02 | -0.42 | > 0.05 |
| nhr-3 | 3.97 | 1.96 | 9.49E-04 | 2.76 | 8.37E-03 |
| nhr-17 | 5.08 | -0.66 | 6.76E-05 | 4.42 | > 0.05 |
| nhr-19 | 1.27 | -0.22 | 7.37E-03 | 0.75 | > 0.05 |
| nhr-21 | 1.29 | -0.17 | 0.03 | 0.53 | > 0.05 |
| nhr-34 | 1.96 | 0.12 | 3.28E-05 | 1.84 | > 0.05 |
| nhr-43 | 3.33 | 1.12 | 4.66E-04 | 2.58 | > 0.05 |
| nhr-45 | 1.46 | -0.70 | 1.31E-05 | 1.72 | > 0.05 |
| nhr-46 | 1.55 | 0.00 | 9.03E-05 | 0.65 | 3.58E-03 |
| nhr-58 | 2.28 | 0.40 | 5.90E-03 | -0.17 | 4.94E-03 |
| nhr-84 | 2.35 | 1.17 | 2.14E-03 | 1.73 | 0.04 |
| nhr-99 | 1.89 | 0.09 | 0.05 | 1.74 | > 0.05 |
| nhr-105 | 3.14 | 0.41 | 1.13E-06 | 1.79 | 2.69E-03 |
| nhr-107 | 1.95 | 0.18 | 0.04 | 0.90 | > 0.05 |
| nhr-112 | 2.40 | 0.13 | 0.01 | 1.07 | > 0.05 |
| nhr-117 | 2.70 | 0.44 | 5.63E-05 | 2.51 | > 0.05 |
| nhr-120 | 3.10 | 1.46 | 1.53E-05 | 1.93 | 3.07E-03 |
| nhr-137 | 2.53 | 0.66 | 8.26E-04 | 2.58 | > 0.05 |
| nhr-156 | 3.28 | 0.87 | 5.01E-03 | 2.18 | > 0.05 |
| nhr-162 | 3.48 | 0.89 | 2.32E-03 | 1.91 | 0.03 |
| nhr-173 | 1.43 | 0.02 | 2.82E-03 | 1.77 | > 0.05 |
| nhr-203 | 1.66 | 0.10 | 0.02 | 1.01 | > 0.05 |
| nhr-284 | 2.88 | 1.06 | 6.96E-04 | 2.00 | > 0.05 |
| nhr-38 | -1.40 | 2.35 | 3.21E-04 | 0.37 | 0.02 |

Supplementary table 6:
| Gene | Function | N2<br>air-500 | pre-33<br>air-500 | P value 1 | pre-33<br>air-4000 | P value 2 |
| --- | --- | --- | --- | --- | --- | --- |
| **Cytoplasm** |  |  |  |  |  |  |
| hsp-12.3 | Small HSP | 2.60 | 0.42 | 0.01 | 0.89 | 0.02 |
| hsp-12.6 | Small HSP | 5.26 | -0.27 | 1.24E-04 | 1.63 | 2.42E-03 |
| hsp-16.1 | Small HSP | 2.44 | -0.48 | 9.32E-05 | 3.92 | 2.86E-03 |
| hsp-16.2 | Small HSP | 4.20 | -1.05 | 2.26E-05 | 4.37 | > 0.05 |
| hsp-16.41 | Small HSP | 2.59 | -0.91 | 2.32E-04 | 3.33 | > 0.05 |
| hsp-16.48 | Small HSP | 2.46 | -0.53 | 2.92E-04 | 3.32 | > 0.05 |
| hsp-17 | Small HSP | 1.99 | -1.29 | 1.81E-04 | 1.02 | > 0.05 |
| F08H9.3 | Small HSP | 6.23 | 3.05 | 0.03 | 3.99 | > 0.05 |
| hsp-70 | HSP70 | 4.53 | 0.82 | 3.59E-04 | 4.05 | > 0.05 |
| F44E5.5 | HSP70 | 3.69 | -1.72 | 1.40E-05 | 3.37 | > 0.05 |
| **Mitochondria** |  |  |  |  |  |  |
| hsp-6 | HSP70 | 1.05 | -0.03 | 0.01 | 0.54 | > 0.05 |
| **Cytoplasm** |  |  |  |  |  |  |
| Msi3p | HSP70 | 0.58 | -1.23 | 5.82E-03 | 0.24 | > 0.05 |

**Supplementary table 7.**
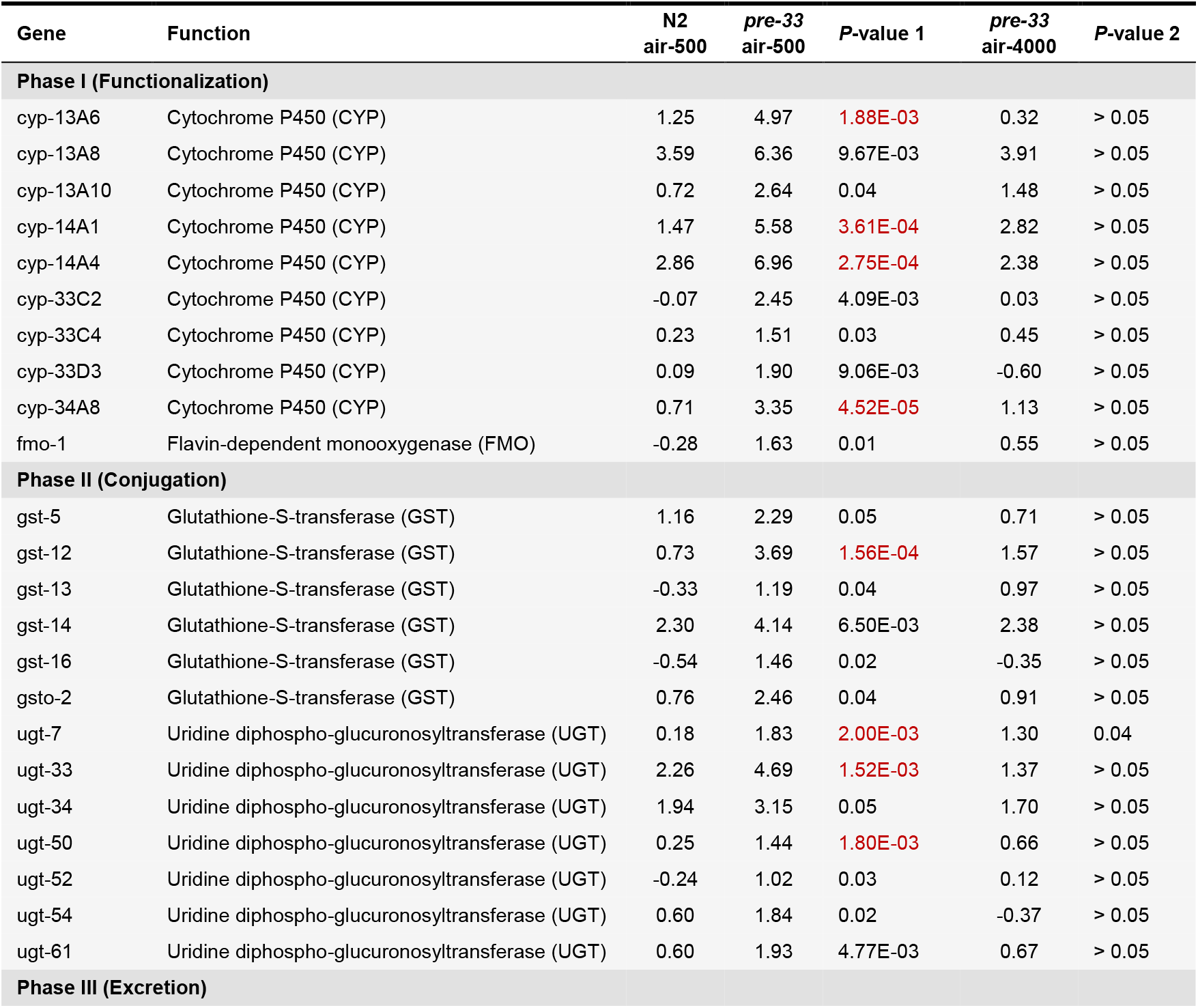
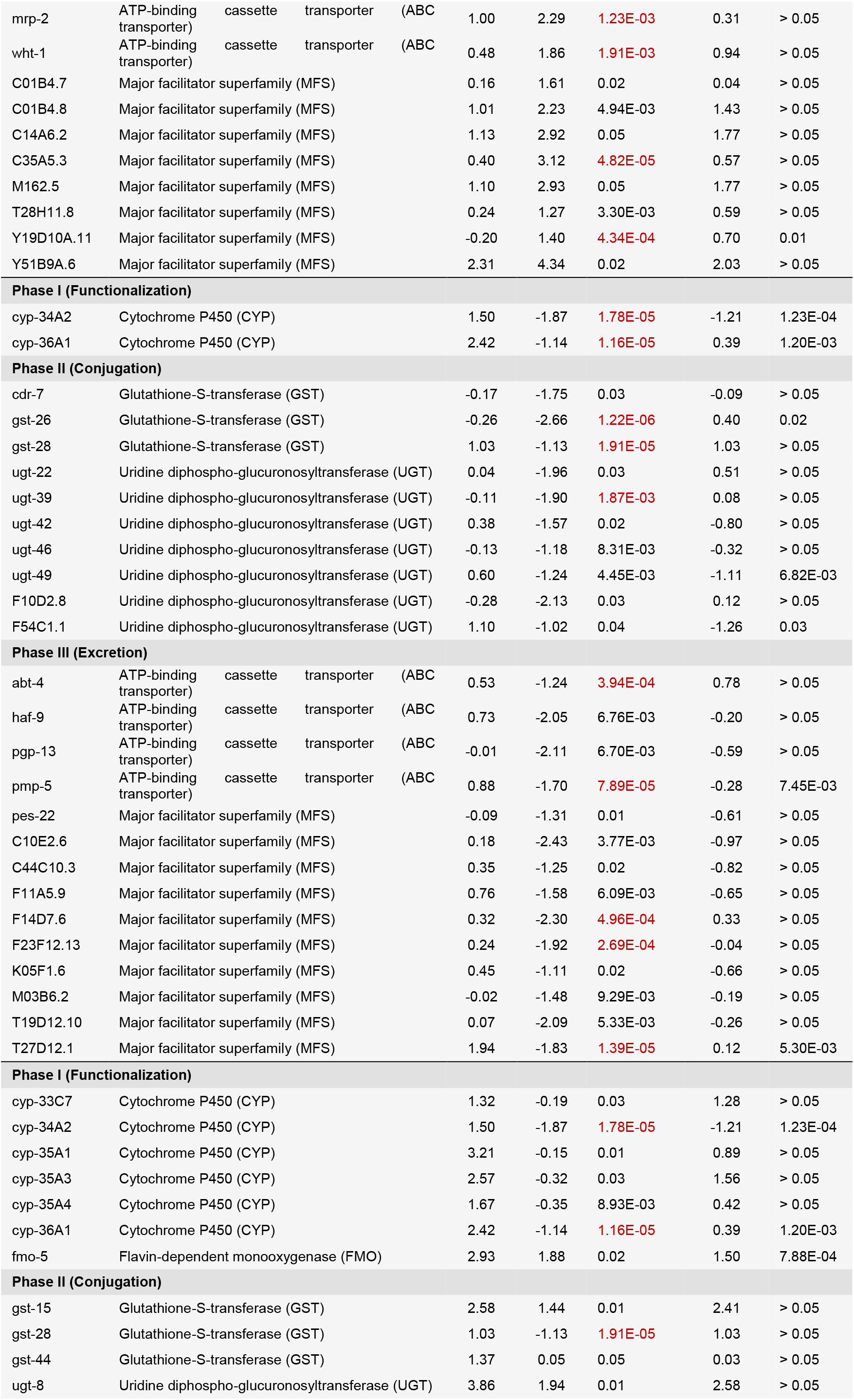
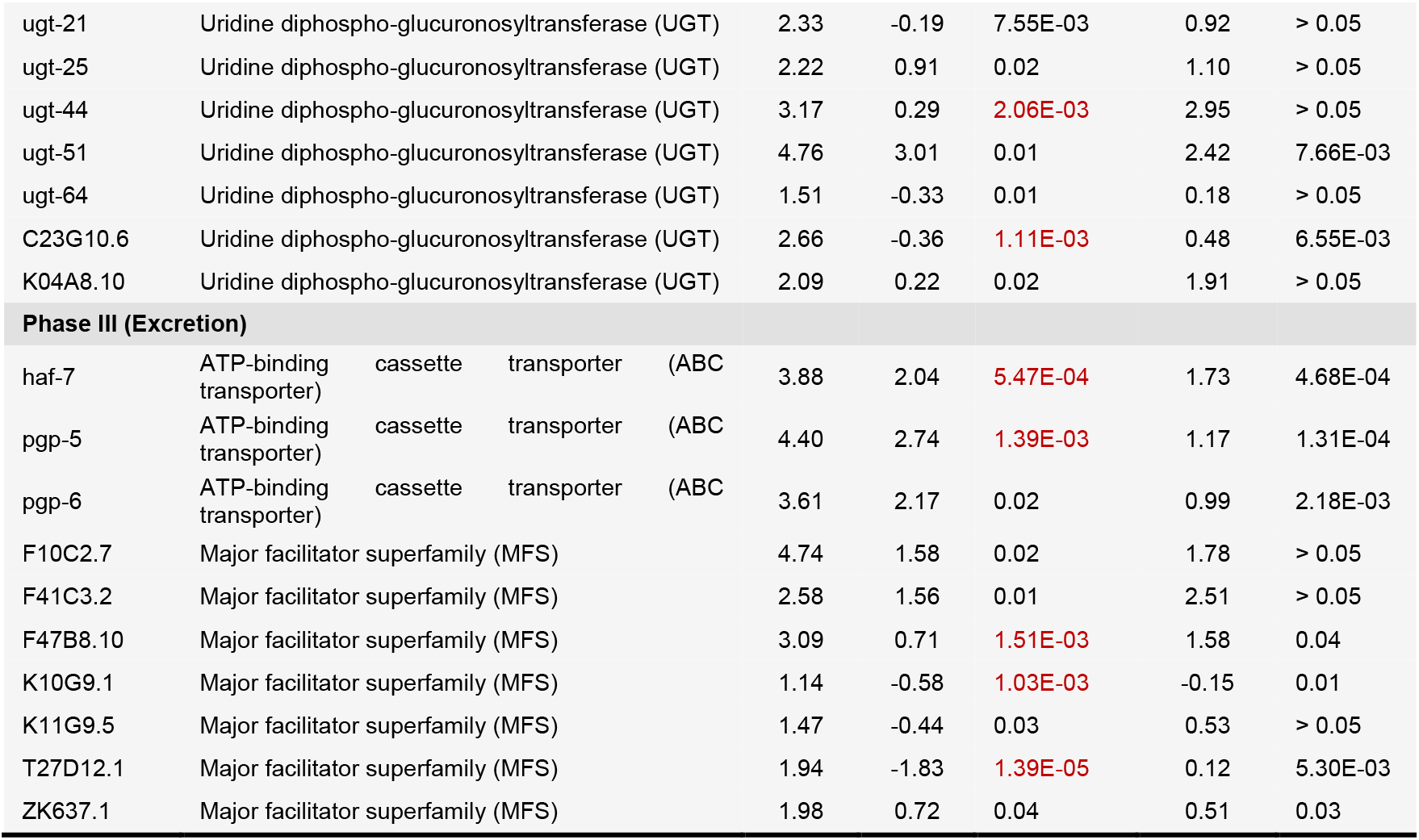

**Supplementary table 8.**

| Gene | Function | Na-Pa | P-value |
| --- | --- | --- | --- |
| bath-1 | BTB and MATH domain containing | 2.62 | 2.35E-03 |
| bath-2 | BTB and MATH domain containing | 2.41 | 1.36E-03 |
| bath-5 | BTB and MATH domain containing | 1.42 | 0.01 |
| bath-7 | BTB and MATH domain containing | 3.33 | 1.19E-03 |
| bath-20 | BTB and MATH domain containing | 1.49 | 0.03 |
| bath-21 | BTB and MATH domain containing | 1.93 | 6.05E-03 |
| bath-22 | BTB and MATH domain containing | 2.11 | 4.81E-04 |
| bath-24 | BTB and MATH domain containing | 1.17 | 9.03E-03 |
| math-3 | MATH domain containing | 4.52 | 2.55E-03 |
| math-4 | MATH domain containing | 3.50 | 2.02E-03 |
| math-10 | MATH domain containing | 4.04 | 0.01 |
| math-14 | MATH domain containing | 3.54 | 8.60E-03 |
| math-15 | MATH domain containing | 3.42 | 2.53E-04 |
| math-20 | MATH domain containing | 4.68 | 1.83E-04 |
| math-21 | MATH domain containing | 4.29 | 2.05E-04 |
| math-22 | MATH domain containing | 3.52 | 5.49E-04 |
| math-26 | MATH domain containing | 3.99 | 5.82E-04 |
| math-27 | MATH domain containing | 4.00 | 9.39E-05 |
| math-39 | MATH domain containing | 3.53 | 4.53E-04 |
| math-40 | MATH domain containing | 3.14 | 7.27E-04 |
| math-41 | MATH domain containing | 2.62 | 1.91E-04 |
| math-50 | MATH domain containing | 4.73 | 9.34E-04 |
| bath-10 | BTB and MATH domain containing | -1.84 | 0.04 |
| bath-16 | BTB and MATH domain containing | -2.56 | 0.01 |
| bath-36 | BTB and MATH domain containing | -1.22 | 0.04 |

**Supplementary table 9:**
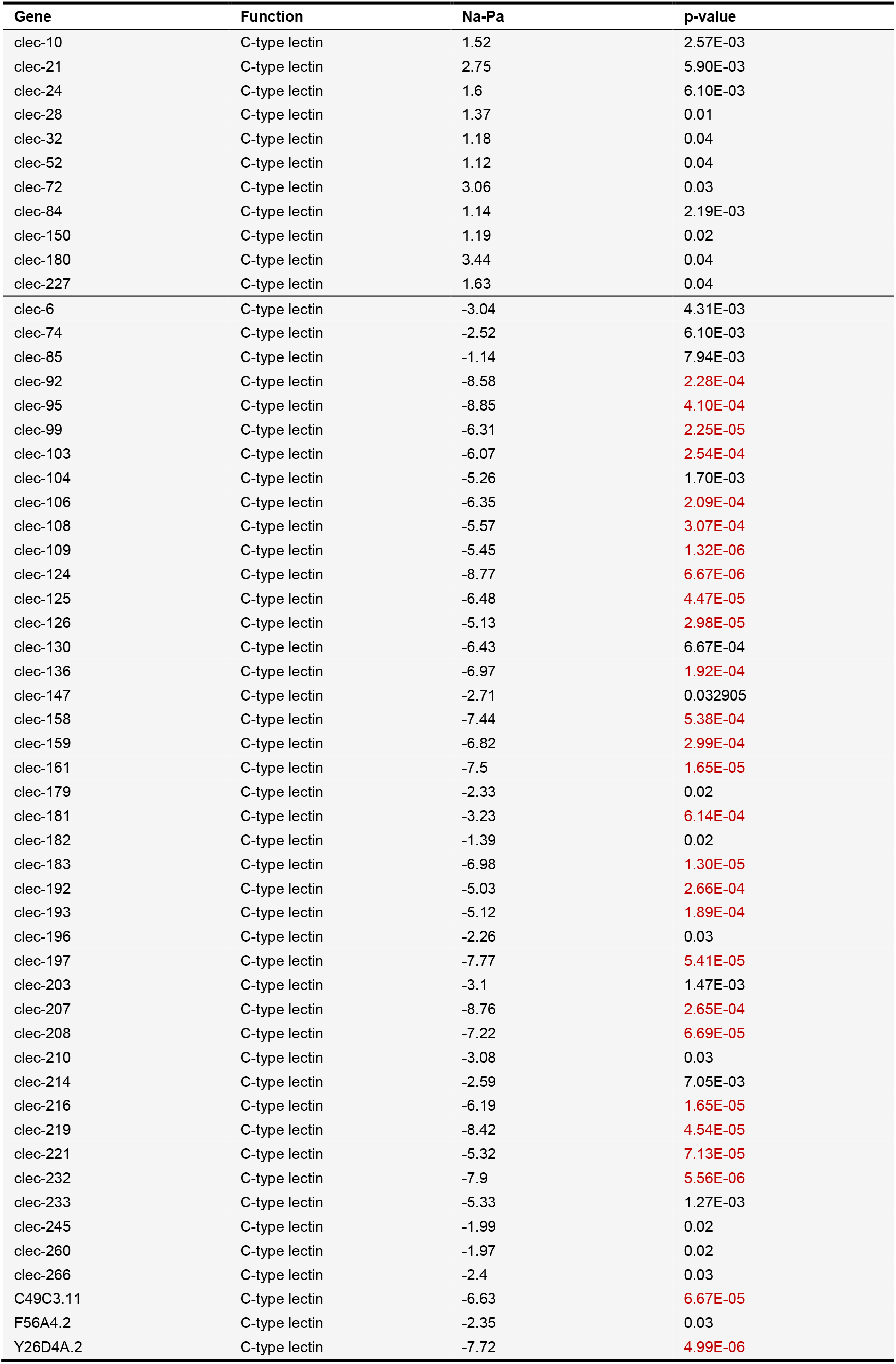

**Supplementary table 10:**

| Gene | Function | Na-Pa | p-value |
| --- | --- | --- | --- |
| F-box A protein |  |  |  |
| fbxa-4 | F-box domain containing | 1.97 | 0.02 |
| fbxa-5 | F-box domain containing | 1.07 | 0.04 |
| fbxa-6 | F-box domain containing | 6.50 | 1.41E-04 |
| fbxa-11 | F-box domain containing | 2.87 | 3.36E-04 |
| fbxa-26 | F-box domain containing | 2.74 | 0.03 |
| fbxa-27 | F-box domain containing | 2.18 | 0.01 |
| fbxa-32 | F-box domain containing | 1.18 | 0.04 |
| fbxa-35 | F-box domain containing | 2.29 | 2.93E-03 |
| fbxa-38 | F-box domain containing | 2.86 | 1.63E-03 |
| fbxa-51 | F-box domain containing | 3.59 | 2.37E-04 |
| fbxa-55 | F-box domain containing | 3.98 | 5.78E-03 |
| fbxa-59 | F-box domain containing | 2.33 | 0.03 |
| fbxa-60 | F-box domain containing | 4.68 | 2.60E-04 |
| fbxa-67 | F-box domain containing | 3.86 | 1.28E-03 |
| fbxa-79 | F-box domain containing | 1.44 | 0.04 |
| fbxa-80 | F-box domain containing | 2.92 | 5.81E-03 |
| fbxa-82 | F-box domain containing | 2.23 | 0.05 |
| fbxa-85 | F-box domain containing | 2.04 | 0.04 |
| fbxa-91 | F-box domain containing | 1.94 | 0.02 |
| fbxa-92 | F-box domain containing | 1.34 | 7.45E-03 |
| fbxa-114 | F-box domain containing | 1.85 | 0.05 |
| fbxa-115 | F-box domain containing | 1.63 | 2.33E-03 |
| fbxa-116 | F-box domain containing | 1.48 | 0.02 |
| fbxa-118 | F-box domain containing | 1.20 | 9.50E-03 |
| fbxa-147 | F-box domain containing | 1.70 | 5.90E-03 |
| fbxa-158 | F-box domain containing | 3.57 | 0.01 |
| fbxa-166 | F-box domain containing | 2.98 | 6.54E-03 |
| fbxa-182 | F-box domain containing | 2.72 | 1.00E-03 |
| fbxa-185 | F-box domain containing | 1.24 | 0.02 |
| fbxa-188 | F-box domain containing | 1.89 | 0.02 |
| fbxa-216 | F-box domain containing | 1.45 | 0.01 |
| F-box A protein |  |  |  |
| fbxa-107 | F-box domain containing | -2.72 | 5.99E-04 |
| fbxa-108 | F-box domain containing | -2.08 | 0.01 |
| fbxa-113 | F-box domain containing | -3.07 | 1.40E-03 |
| fbxa-129 | F-box domain containing | -1.61 | 8.14E-03 |
| fbxa-191 | F-box domain containing | -4.20 | 6.09E-04 |
| fbxa-192 | F-box domain containing | -2.67 | 5.55E-03 |
| fbxa-201 | F-box domain containing | -1.08 | 0.01 |
| fbxa-217 | F-box domain containing | -3.33 | 3.44E-04 |
| F-box B protein |  |  |  |
| fbxb-36 | F-box domain containing | -1.49 | 0.02 |
| fbxb-56 | F-box domain containing | -1.80 | 5.62E-04 |
| fbxb-75 | F-box domain containing | -2.13 | 0.02 |
| F-box C protein |  |  |  |
| fbxc-36 | F-box domain containing | -1.72 | 0.02 |
| fbxc-48 | F-box domain containing | -1.24 | 0.02 |
| fbxc-49 | F-box domain containing | -2.19 | 4.41E-05 |
| fbxc-51 | F-box domain containing | -2.21 | 0.05 |

**Supplementary table 11:**

| Gene | Function | Na-Pa | p-value |
| --- | --- | --- | --- |
| clec-10 | C-type lectin | 1.52 | 2.57E-03 |
| clec-21 | C-type lectin | 2.75 | 5.90E-03 |
| clec-24 | C-type lectin | 1.6 | 6.10E-03 |
| clec-28 | C-type lectin | 1.37 | 0.01 |
| clec-32 | C-type lectin | 1.18 | 0.04 |
| clec-52 | C-type lectin | 1.12 | 0.04 |
| clec-72 | C-type lectin | 3.06 | 0.03 |
| clec-84 | C-type lectin | 1.14 | 2.19E-03 |
| clec-150 | C-type lectin | 1.19 | 0.02 |
| clec-180 | C-type lectin | 3.44 | 0.04 |
| clec-227 | C-type lectin | 1.63 | 0.04 |
| clec-6 | C-type lectin | -3.04 | 4.31E-03 |
| clec-74 | C-type lectin | -2.52 | 6.10E-03 |
| clec-85 | C-type lectin | -1.14 | 7.94E-03 |
| clec-92 | C-type lectin | -8.58 | 2.28E-04 |
| clec-95 | C-type lectin | -8.85 | 4.10E-04 |
| clec-99 | C-type lectin | -6.31 | 2.25E-05 |
| clec-103 | C-type lectin | -6.07 | 2.54E-04 |
| clec-104 | C-type lectin | -5.26 | 1.70E-03 |
| clec-106 | C-type lectin | -6.35 | 2.09E-04 |
| clec-108 | C-type lectin | -5.57 | 3.07E-04 |
| clec-109 | C-type lectin | -5.45 | 1.32E-06 |
| clec-124 | C-type lectin | -8.77 | 6.67E-06 |
| clec-125 | C-type lectin | -6.48 | 4.47E-05 |
| clec-126 | C-type lectin | -5.13 | 2.98E-05 |
| clec-130 | C-type lectin | -6.43 | 6.67E-04 |
| clec-136 | C-type lectin | -6.97 | 1.92E-04 |
| clec-147 | C-type lectin | -2.71 | 0.032905 |
| clec-158 | C-type lectin | -7.44 | 5.38E-04 |
| clec-159 | C-type lectin | -6.82 | 2.99E-04 |
| clec-161 | C-type lectin | -7.5 | 1.65E-05 |
| clec-179 | C-type lectin | -2.33 | 0.02 |
| clec-181 | C-type lectin | -3.23 | 6.14E-04 |
| clec-182 | C-type lectin | -1.39 | 0.02 |
| clec-183 | C-type lectin | -6.98 | 1.30E-05 |
| clec-192 | C-type lectin | -5.03 | 2.66E-04 |
| clec-193 | C-type lectin | -5.12 | 1.89E-04 |
| clec-196 | C-type lectin | -2.26 | 0.03 |
| clec-197 | C-type lectin | -7.77 | 5.41E-05 |
| clec-203 | C-type lectin | -3.1 | 1.47E-03 |
| clec-207 | C-type lectin | -8.76 | 2.65E-04 |
| clec-208 | C-type lectin | -7.22 | 6.69E-05 |
| clec-210 | C-type lectin | -3.08 | 0.03 |
| clec-214 | C-type lectin | -2.59 | 7.05E-03 |
| clec-216 | C-type lectin | -6.19 | 1.65E-05 |
| clec-219 | C-type lectin | -8.42 | 4.54E-05 |
| clec-221 | C-type lectin | -5.32 | 7.13E-05 |
| clec-232 | C-type lectin | -7.9 | 5.56E-06 |
| clec-233 | C-type lectin | -5.33 | 1.27E-03 |
| clec-245 | C-type lectin | -1.99 | 0.02 |
| clec-260 | C-type lectin | -1.97 | 0.02 |
| clec-266 | C-type lectin | -2.4 | 0.03 |
| C49C3.11 | C-type lectin | -6.63 | 6.67E-05 |
| F56A4.2 | C-type lectin | -2.35 | 0.03 |
| Y26D4A.2 | C-type lectin | -7.72 | 4.99E-06 |
| pqn-84 | Galectin | -5.74 | 3.10E-04 |
| F40H6.5 | Galectin | -1.49 | 2.89E-04 |
| F46A8.3 | Galectin | -7.18 | 1.37E-06 |
| F49F1.9 | Galectin | -7.41 | 1.09E-05 |
| F49F1.10 | Galectin | -4.82 | 1.21E-04 |
| F49F1.11 | Galectin | -7.04 | 1.28E-05 |

